# Low-Noise Finite-Bandgap FET Biosensors in the Nonlinear Gouy-Chapman-Stern Region: A Universal Calibration Curve

**DOI:** 10.64898/2026.09.16.752103

**Authors:** Feng Gao, Tyler Moorman, Tiger Haoran Shi, Sizhe Ma, Kai Ni, Satyajyoti Senapati, Hsueh-Chia Chang

## Abstract

Finite-bandgap charge-based FET biosensors are known to be insensitive to electronic tunneling noise in the semiconductor and interference from weakly-charged proteins in physiological samples. However, their signal is still corrupted by pH fluctuations and condensing counterions in the bulk electrolyte. Here, we demonstrate that electrolyte noise is screened by the Stern layer beyond the linear Debye-Hückel limit, when the interfacial potential drop is larger than the thermal voltage. Using a fully nonlinear Gouy-Chapman-Stern (GCS) theory, a universal calibration curve is derived and shown to collapse the low-noise FET signals for miRNA, extracellular vesicles (EV), and protein, captured either directly or through silica and magnetic nanoparticle charge reporters, over eleven decades of analyte concentration. The theory involves the dissociation constant *K_D_* and a gain parameter χ, which represents the competing effects of the reporter charge and buffer Debye screening that must be large to produce the low-noise GCS condition. The limit of detection (LOD) is shown to be *K_D_*/χ, which can be optimized by varying the probe density, charge reporter, and buffer solution. An optimized hybridization buffer then yields χ ∼ 22 (LOD ∼1 pM) for miRNA detection, and multivalent capture reduces *K_D_* by four orders of magnitude for EV detection (relative to protein detection) with χ ∼ 34 (LOD ∼10 fM).

---

Electrochemical biosensors that convert a molecular recognition event into an electrical current signal have become central to modern bioanalysis, particularly for the nascent field of liquid biopsy, yet quantification of what binds the probes at the sensor surface and what is measured at its output remains highly empirical. Almost universally, each combination of target, receptor, and measurement condition is characterized by its own calibration curve, obtained by fitting a response to a set of standards and used only within the narrow window in which it was acquired.^1^ The underlying mechanism responsible for such non-specificity is the myriad of redox charge-transfer reactions with kinetics sensitive to pH and interfering redox agents. A signal recorded for one analyte cannot readily be compared to that of another, and a calibration established in one buffer rarely transfers to a second. Consequently, tedious calibration, sometimes for each sample, is required for quantitative electrochemical sensing. The field has produced steadily more sensitive and more miniaturized devices, but a quantitative, science-based account of transduction, which allows optimization of the detection buffer, has remained comparatively underdeveloped.^2,3^

Field-effect transistor (FET) biosensors are an attractive alternative to address this issue. Without charge transfer from the reporters or analytes, FET charge transduction is less sensitive to interfering redox agents in the buffer. As the Zeta potential of proteins, the most abundant interfering agents in physiological samples, is typically less than 20 mV^4^, it is also insensitive to non-specific adsorption of proteins onto the sensor. Moreover, since how charge near a gate electrode modulates the channel potential (and therefore the channel current within the transistor) is well-known from classical semiconductor physics, the output can be related to the charge with a universal, scientifically derived and validated correlation rather than case-by-case empirical fitting. Since the first ion-sensitive devices, this lineage has advanced dramatically in sensitivity.^5^ Channels based on silicon nanowires^6^ and, more recently, on graphene^7,8^, MXene^9,10^, and other two-dimensional materials^11^ have achieved sub-femtomolar and even single-molecule detection. These advances are impressive but come at a practical cost: such devices rely on materials and fabrication processes that are expensive, difficult to reproduce at scale, and poorly suited to disposable, multiplexed formats, which has limited their translation beyond specialized laboratories. The extended-gate FET (EGFET) offers a pragmatic alternative. It separates a disposable, chemically functionalized sensing electrode from a reusable, commercial transistor, is compatible with low-cost planar fabrication, and scales readily into arrays.^12–14^ This separation also allows independent optimization of the semiconductor transducer, which is reusable, and the incubation buffer/probe at the sensor electrode.

FET charge sensors, however, still suffer from some reproducibility issues. In a zero-bandgap Dirac semiconductor such as 2D graphene, a small interfacial charge produces a large current change, but that current carries a correspondingly large 1/f noise generally attributed to the trapping and release of carriers by tunneling, which makes such devices as difficult to calibrate as electrochemical sensors. Their practical detection limits are well above the concentrations at which a response is already visible.^15^ Suppressing that noise requires costly measures, magnetic Hall-effect readout or AC cycling of the gate about the Dirac point^16^, and the resulting picomolar limits for nucleic acids remain comparable to far less expensive electrokinetic charge sensors.^17,18^ A finite-bandgap transistor operated below threshold avoids the problem at its source: tunneling is minimized, and the drain current depends exponentially rather than linearly on the gate potential^19^, so the sensitivity lost because the gating field does not penetrate a three-dimensional semiconductor channel as deeply as a two-dimensional one is recovered by the exponential amplification. The finite bandgap is also what makes the extended-gate format of EGFET possible, since a two-dimensional sheet cannot be contacted directly by an electrode and therefore requires the sample to sit on the transducer itself, precluding both reuse and disposable large arrays.

There are two remaining obstacles that have kept EGFET biosensing from being widely applied. The first relates to Debye screening: charge-based transduction senses only the charge lying within the Debye screening length on the electrode sensor.^7,20^ This implies that the signal varies with the ionic strength, although this variation can be corrected with classical electrokinetic theory.^21^ More importantly, for solutions of physiological ionic strength, the Debye length collapses to well below a nanometer, so much of a captured analyte’s charge is screened.^20,22^ A low-ionic strength buffer is hence preferred. However, low-ionic strength buffers often prevent nucleic acid hybridization^23^ and cause lysis of extracellular vesicles.^24^ An optimum ionic strength may hence exist but this remains unexplored because a theory that relates the analyte charge to the EGFET signal remains unexplored due to an irreproducibility issue discussed below.

The second and more crucial obstacle is counterion condensation or pH fluctuation that can alter the charge or even invert it.^25,26^ Such noise-driven surface charge variation often produces widely varying reporter signals in FET or any charge-based sensor technology. Quite often, calibration curves with extremely low signal-to-noise ratio or with nearly discontinuous jumps are reported.^12,18, 19^ As the tunneling noise in the semiconductor has been suppressed, this electrolyte noise becomes the principal obstacle to quantification by or optimization of a finite-bandgap EGFET sensor.

There are, in fact, buffer/reporter conditions that can mitigate this variability due to bulk electrolyte fluctuations. It has been shown that, for highly charged surfaces, with Zeta potential higher than *RT/F* (∼26 mV at room temperature), a Stern layer of well-packed hydrated counterions exists on the charged surface.^21^ These Stern-layer hydrated ions and the ordered water molecules around them do not change the charge on the surface and, in fact, prevent condensation of other counterions.^28^ They electrically insulate the Stern layer so that it is not sensitive to the bulk electrolyte conditions. Measurements of membrane and nanoslot conductance have also shown that they are insensitive to the bulk ionic strength when the Donnan potential exceeds *RT/F*.^29–31^ It is reasonable to assume that, with a sufficiently high surface density of charged analytes or reporters on the electrode sensor of an EGFET, a potential drop higher than *RT/F* and a layer of concentrated hydrated counterions will also appear on the nucleic acids to screen other counterions.^32^ Most proteins and extracellular vesicles (Zeta potential of <20 mV even at low ionic strengths) are too weakly charged to form this protective Stern layer. Their signal is hence weak with large variations. A more highly charged reporter would then need to be added, as we have done for the electrokinetic charge sensor.^33^

If it is true that a high effective Zeta potential on the sensor electrode due to the analyte or its reporter can suppress noise level, it would be advantageous to use high-density probes on the EGFET sensor electrode. High-probe density also has another advantage—multivalent capture. Extracellular vesicles (EVs) are larger than 30 nm, and hence if the antibody probes are less than 30 nm apart, which is quite feasible with most functionalization protocols, there could be multiple antibody-antigen bindings for each EV with multiple antigens. Binding by multivalent antibodies, sometimes referred to as avidity, has been shown to produce a reduction in the dissociation constant *K_D_* by several orders of magnitude in immunocapture.^34,35^ Other than an orders-of-magnitude reduction in *K_D_*, multivalent antibodies have been shown to reduce the dissociation rate constant *k_off_* by the same magnitude.^36^ Hence, selectivity for target capture can also be improved even if the competing interfering agent is much more abundant. For EVs, this advantage of multi-valent capture for enhancing both sensitivity and selectivity can be realized just by increasing the probe density, without special multi-valent antibodies.

Here we validate low-noise EGFET sensing in the GCS (Gouy-Chapman-Stern) region, with an effective Zeta potential larger than *RT/F*, and, using GCS theory, derive a universal continuous calibration curve for all analytes that allows accurate quantification and easy optimization. This calibration curve allows us to optimize the detection buffer and multi-valent capture strategies. We construct a 36-well EGFET array in which all sensing electrodes share a single transistor and are read under identical device conditions, so that entire concentration series and multiple ionic strengths can be acquired in parallel without transducer-to-transducer variation. To bring weakly charged targets within reach of a charge-based readout, we introduce a charged nanoparticle (NP) reporter, so that binding of the analyte-reporter complex to the electrode sensor delivers a well-defined charge to the gate. Using this platform, we measure three chemically distinct classes of liquid biopsy targets across a range of ionic strengths: synthetic miR-9 and miR-21 as small, negatively charged nucleic acids detected directly, purified EVs detected through their charge reporters, and the protein endothelin-1 (EDN1) detected by the same binding reporter. The responses of these most common liquid biopsy targets^37^ are collapsed by a universal calibration curve derived from electrolyte theory, with an explicit limit of detection (LOD) that allows buffer and probe density optimization.

## EXPERIMENTAL SECTION

Chemical reagents and materials are listed in the Supporting Information.

### Extended Gate Fabrication

A polycarbonate (PC) sheet is used as the substrate for gold deposition to form the extended gate. A thin plastic mask bearing the designed electrode pattern was cut and applied to the substrate using a cutter plotter. Prior to metal deposition, the exposed substrate surface was rinsed with isopropyl alcohol (IPA) and treated with plasma to remove organic residues and improve adhesion. A 5 nm titanium adhesion layer was then deposited, followed by 50 nm of gold at a rate of 1.0 Å/s, using an Angstrom Engineering evaporator (Cambridge, Ontario, Canada). The gold-coated substrate was immersed in 2 mM 11-MUA in 200-proof ethanol for 12 hours at room temperature to form a carboxyl-terminated self-assembled monolayer (SAM). The substrate was then rinsed with 200-proof ethanol under gentle sonication to remove excess 11-MUA. Carboxyl groups were activated via EDC/NHS chemistry. A solution of 25 mM Sulfo-NHS and 100 mM EDC in 0.1 M MES buffer (pH 4.7) was applied to each gold sensing pad for 30 min, followed by rinsing with 1× DPBS. Next, 0.1 mg/mL antibody or 1 µM probe miRNA in 1× DPBS was dispensed onto each pad and incubated for 24 hours at 4 °C to complete the reaction. Finally, self-designed 3D-printed wells were sealed onto the substrate to form the 36-well array extended gate.

### Charge Reporter Functionalization

Charge reporters were prepared by conjugating a reporter antibody to the carboxylated silica NPs by EDC/NHS chemistry. The same procedure was used for the carboxylated magnetic NPs. A 250 µL aliquot of the silica particles from stock was mixed with 750 µL of 0.1 M MES buffer in a low-binding Eppendorf tube, followed by vortexing, and centrifugation at 15,000 × g for 10 min to pellet the particles. The supernatant was removed. The pellet was resuspended by sonication in 1 mL of 100 mM EDC and 25 mM Sulfo-NHS in 0.1 M MES buffer and incubated for 30 min on an orbital shaker. The particles were centrifuged for 10 min, the supernatant was removed, and the pellet was redispersed by sonication in 900 µL of 1× DPBS. 100 µL of 0.1 mg/mL reporter antibody was added, and the suspension was vortexed and incubated on a rotator at 4 °C for 24 h.

### Electrical Characterization of Sensing Signal by the Extended-Gate Field-Effect Transistor

Prior to electrical measurement, miRNA, EV, or EDN1 protein samples were incubated in each sensor well for 60 min in specific ionic strengths, followed by 4X DPBS wash to remove non-specific binding and then changed back to the same ionic strength as incubation. miRNA targets were measured from 1 × 10^−15^ to 1 × 10^−5^ M in 0.15 to 150 mM buffer, EVs from 7.64 × 10^−16^ to 7.64 × 10^−11^ M and EDN1 protein from 2 × 10^−12^ to 2 × 10^−7^ M in 1.5 mM and 150 mM. Each concentration was measured in three wells (n = 3). The electrical characteristics of the fabricated biosensor, specifically the transfer characteristic between drain current I_DS_ and gate voltage V_GS_ was measured using Keithley 4200A (Keithley Instruments, Cleveland, OH, USA). The extended-gate configuration was implemented by using a commercial n-type enhancement-mode MOSFET as the transducer and the extended Au electrode as the sensing film. The gate was extended by connecting the MOSFET gate terminal to the electrode (Figure 1b). The gate potential was applied through an Ag/AgCl reference electrode (DRI-REF, World Precision Instruments, Sarasota, FL) immersed in each well, which contained 1× (150 mM), 0.1× (15 mM), 0.01× (1.5 mM), or 0.001× (0.15 mM) DPBS, or deionized water for the salt-free condition. The gate voltage was swept linearly from −1.5 to 2 V while the drain voltage was held at 0.1 V and the source terminal was grounded. I_DS_ was recorded throughout the sweep.

### Isolation of Extracellular Vesicle

EVs were isolated from DiFi conditioned medium by differential ultracentrifugation. The medium was first concentrated from 10 mL to 1 mL on a 100 kDa filter. The concentrate was centrifuged at 12,000 × g for 20 min to remove large particles, and the supernatant was passed through a 220 nm filter to remove residual debris. The filtrate was layered over 3 mL of DPBS in a 4 mL ultracentrifuge tube and ultracentrifuged at 167,000 × g for 1.5 h in a swinging-bucket rotor (Beckman Coulter SW60Ti), which pellets vesicles more efficiently than a fixed-angle rotor. The supernatant was removed down to a residual volume of about 0.2 mL to avoid disturbing the pellet, and the pellet was resuspended in 0.5 mL of DPBS. The suspension was layered over 15.5 mL of DPBS in a 17 mL ultracentrifuge tube and ultracentrifuged at 167,000 × g for 4.5 h (Beckman Coulter SW32.1Ti). The final pellet was resuspended in DPBS and passed through a 300 kDa filter to remove soluble protein.

### Nanoparticle Tracking Analysis

Particle concentration of purified EVs was determined by nanoparticle tracking analysis (NTA) using a NanoSight NS300 (Malvern Panalytical, Malvern, UK) housed at the Harper Cancer Research Institute, University of Notre Dame, following the NanoSight NS300 user manual. Each sample was analyzed in triplicate using a serial dilution series in 1× DPBS to ensure measurement reliability. Camera settings were as follows: capture gain 8, camera level 10, focus 450, process gain 10, and detection threshold 3. Final particle concentrations were scaled to reflect the original, undiluted sample volumes. Particle counts per mL were converted to molar concentrations. The 1000-fold diluted stock gave 4.60 × 10^7^ particles per mL; the undiluted preparation contained 4.60 × 10^10^ particles per mL, or 7.64 × 10^−11^ M.

### Zeta Potential Measurement

We measured the zeta potential of purified EVs, EDN1 protein, miR-9, carboxylated silica NPs, and carboxylated magnetic NPs using a Malvern Zetasizer Wave II (Microtrac). All samples were dispensed in 1× DPBS to maintain consistent salt concentration.

### Curve Fitting

For each condition, the signal S = log_10_(I_B_/I_T_) from each of the three replicate wells was transformed as Y = sinh(αS), with α = n ln(10)/2 and the subthreshold ideality factor fixed at n = 1.83 from the transfer characteristic. The transformed replicates were fitted to Y = χ[A]/(K_D_ + [A]), the sinh-transformed form of equation (6), by nonlinear least squares with χ and K_D_ as the only free parameters, both constrained to positive values. The Debye length λ_D_ was computed from the buffer ionic strength and held fixed, and λ_S_ was obtained as λ_D_/χ.

Uncertainties were estimated by resampling the three replicates at each concentration with replacement and refitting, over 2,000 bootstrap datasets. Table 1 reports each fitted or derived estimate ± one standard deviation of its bootstrap distribution. Signal-to-noise ratios were calculated from the untransformed signals as the mean S divided by the standard deviation of the three wells at each concentration, with S/N = 3.3 taken as the detection threshold. The miR-9 series without added salt and the reporter-free EV and EDN1 controls showed no concentration dependence and did not yield identifiable binding constants. For EDN1, Table 1 reports the monoclonal-capture/polyclonal-reporter scheme only.

## RESULTS AND DISCUSSION

### Charge-Based Signal Transduction and the 36-Well EGFET Array

The EGFET biosensor converts a molecular binding event into an electrical signal through the chain of charge-mediated steps in Figure 1a. Target capture at the probe-functionalized Au gate raises the fractional surface occupancy θ. Because each bound target carries net charge, this changes the interfacial charge density Δσ, which shifts the effective threshold voltage of the extended gate for the transistor by ΔV_analyte_. The shift is read out as the log-ratio of the baseline and target drain currents, log_10_(I_B_/I_T_). The premise underlying every link is that the captured species must be charged. The transducer responds to charge delivered to the gate, not to mass or to binding itself.

**Figure 1.**
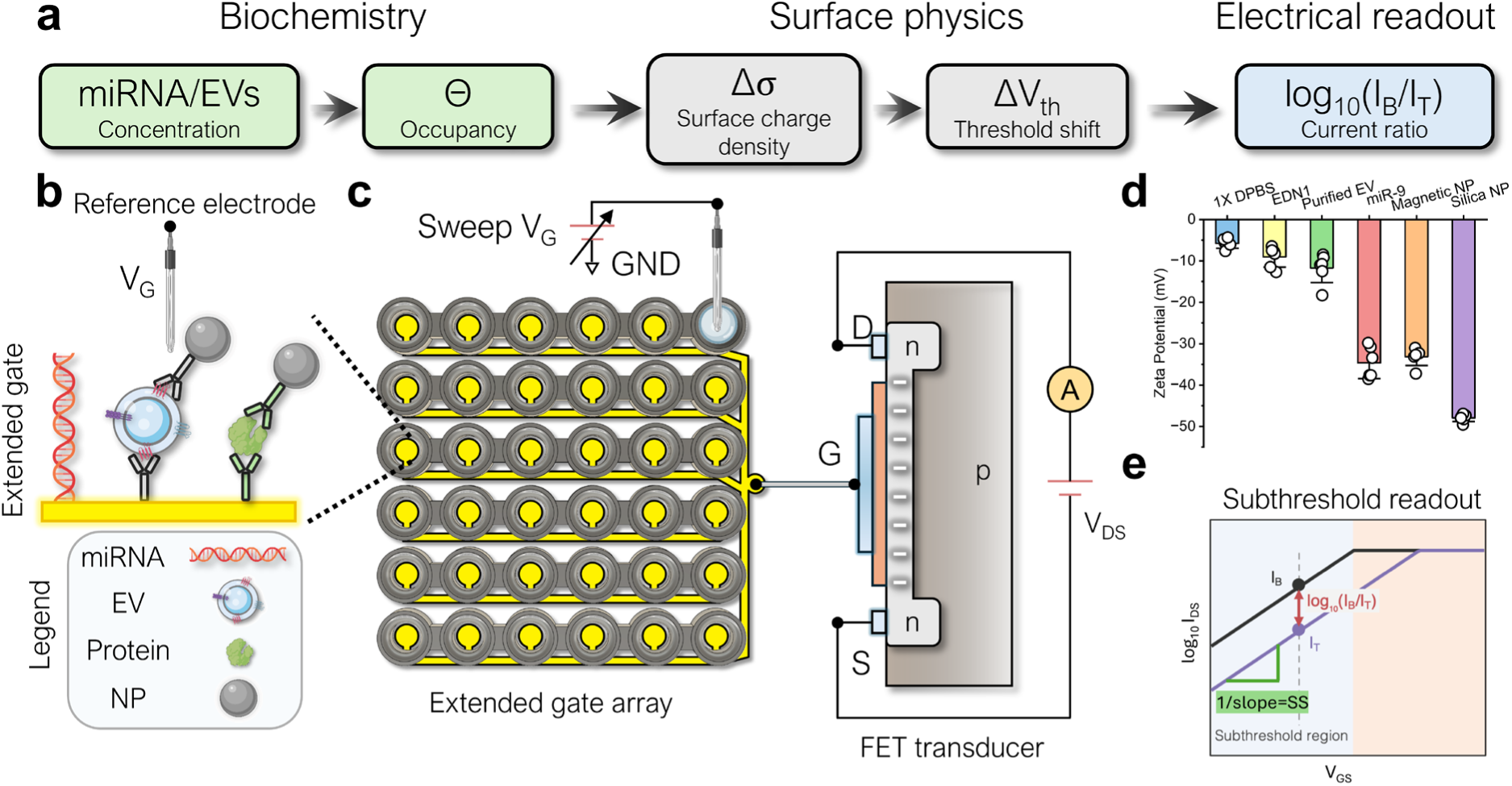
Charge-based signal transduction on the 36-well EGFET array. (a) Signal transduction chain linking biochemistry, surface physics, and electrical readout. (b) Functionalized extended-gate surface, showing direct hybridization of miRNA and antibody-mediated capture of EVs and proteins with highly charged NP reporters. (c) The 36-well extended-gate array, in which gold electrodes patterned on a polycarbonate substrate are tied to the gate terminal G of a single shared n-type MOSFET transducer. The gate voltage VG is applied through an Ag/AgCl reference electrode in each well while the drain current is recorded at fixed V_DS_. (d) Zeta potential of 1× DPBS, EDN1 protein, purified EVs, miR-9, magnetic NPs, and silica NPs measured in 1× DPBS. (e) Subthreshold readout: the baseline current I_B_ and target current I_T_ are read at the same V_GS_ within the subthreshold region, where the inverse slope of the semi-log transfer curve defines the subthreshold swing SS.

Figure 1b shows the functionalized extended gate that presents these binding events to the transducer. A carboxyl-terminated 11-MUA self-assembled monolayer on the gold electrode is activated by EDC/NHS chemistry and coupled either to a single-stranded DNA probe or to a capture antibody, so that one common surface chemistry supports chemically distinct targets. Small nucleic acid targets are detected directly: miRNA hybridizes to the immobilized probe and brings its charge phosphate backbone within the Debye screening length of the gate. Larger and more weakly charged targets, EVs and proteins, are instead captured by their respective antibodies and subsequently labeled with a highly charged NP reporter, so that the charge delivered to the gate is set by the reporter rather than by the target itself. In all three types of analytes, the gate potential is applied through an Ag/AgCl reference electrode immersed in the same well, so that the surface potential established by the captured layer is referenced to the bulk electrolyte.

The sensing surface is arranged as a 36-well array as shown in Figure 1c and the optical photo in Figure S1. Gold electrodes patterned on a polycarbonate substrate each serve as an independent extended gate, and 3D-printed wells define 36 isolated reaction chambers on the same chip. The Au electrodes are tied together by a common connector that routes to the gate terminal of a single shared n-type MOSFET transducer, while an Ag/AgCl reference electrode is inserted into each well to apply the gate voltage V_GS_. During measurement, the drain-source voltage V_DS_ is held constant while the drain current I_DS_ is recorded. This shared-transducer design allows a full concentration series, or several buffer conditions in parallel, to be acquired in a single run under identical device conditions, so that every curve in a family is measured against the same transistor. Because all 36 electrodes share one gate connection, we verified that the solution in one well does not perturb the reading at another. Changing neighboring wells to 1× PBS or to 10% HCl left the transfer curve at the monitored well unchanged, whereas changing the monitored well buffer shifted it (Figure S3).

To confirm that the target and reporter species carry the charge the transduction scheme requires, we measured the Zeta potential of each analyte and charge reporter (Figure 1d). The 1× DPBS blank sat near zero, at −5.78 mV. EDN1 protein and purified EVs were both weakly charged, at −9.05 and −11.71 mV, well below the thermal voltage of 25.9 mV. In contrast, miR-9 was strongly negative at −34.62 mV, consistent with its phosphate backbone, and is therefore suited to direct detection. The magnetic and silica NP reporters were −33.15 and −47.90 mV. Only hybridized miRNA on the sensor electrode can, at high occupancy, produce a net Zeta potential that exceeds the thermal voltage *RT/F*. Reporters are therefore necessary for EDN1 and EVs.

Finally, Figure 1e shows how a change at the interface is converted into the reported signal. The drain current is recorded as log_10_(I_DS_) against V_GS_ before and after target incubation, and the two curves are compared at a single gate voltage within the subthreshold region, where log_10_(I_DS_) is linear in V_GS_. Because the captured targets and reporters are negatively charged, binding raises the effective threshold voltage and translates the curve toward higher V_GS_, so that at a fixed operating point the current falls from its baseline value I_B_ to the target value I_T_. The vertical separation between the two curves, log_10_(I_B_/I_T_), is therefore a direct measure of the binding-induced shift in interfacial potential, and it is this quantity that we report throughout. The inverse slope of the linear portion defines the subthreshold swing SS, which sets the conversion between potential shift and log-current change and is a property of the transistor alone, not of the surface chemistry or the buffer. Since a single shared transducer serves all 36 wells, this conversion factor is common to every analyte and ionic strength measured on the array.

### Transfer Characteristics and Concentration Response

Figure 2 shows the measured transfer characteristics, i.e., the drain current I_DS_ as a function of gate voltage V_GS_, for miR-9 across four buffer ionic strengths, with complementary target strand concentration from 1 fM to 10 µM recorded in 150 mM, 15 mM, 1.5 mM, and 0.15 mM ionic strength buffer DPBS. In every buffer, the curves shift systematically with concentration, but the separation between successive concentrations does not widen monotonically with dilution: it is widest in 1.5 mM, while the 15 mM and 0.15 mM panels are comparable to one another. Reduced screening accounts for the initial gain, since a lower salt concentration increases the Debye length, so a larger fraction of each bound target’s charge is sensed at the gate and a given amount of binding produces a larger threshold shift. Screening alone would predict the widest spacing at the lowest ionic strength, so the reversal below 1.5 mM indicates a competing loss: at very low ionic strength the duplex is destabilized, and the electrode baseline is less stable, offsetting the longer screening length. An optimum ionic strength therefore exists for a given miRNA probe–target pair. Across all four buffers, the curves are translated along the voltage axis without any change in semi-log slope, so binding enters the readout only through the effective threshold voltage while the subthreshold swing remains a property of the transistor. The same conversion between interfacial potential shift and log-current change therefore applies at every concentration and in every buffer.

**Figure 2.**
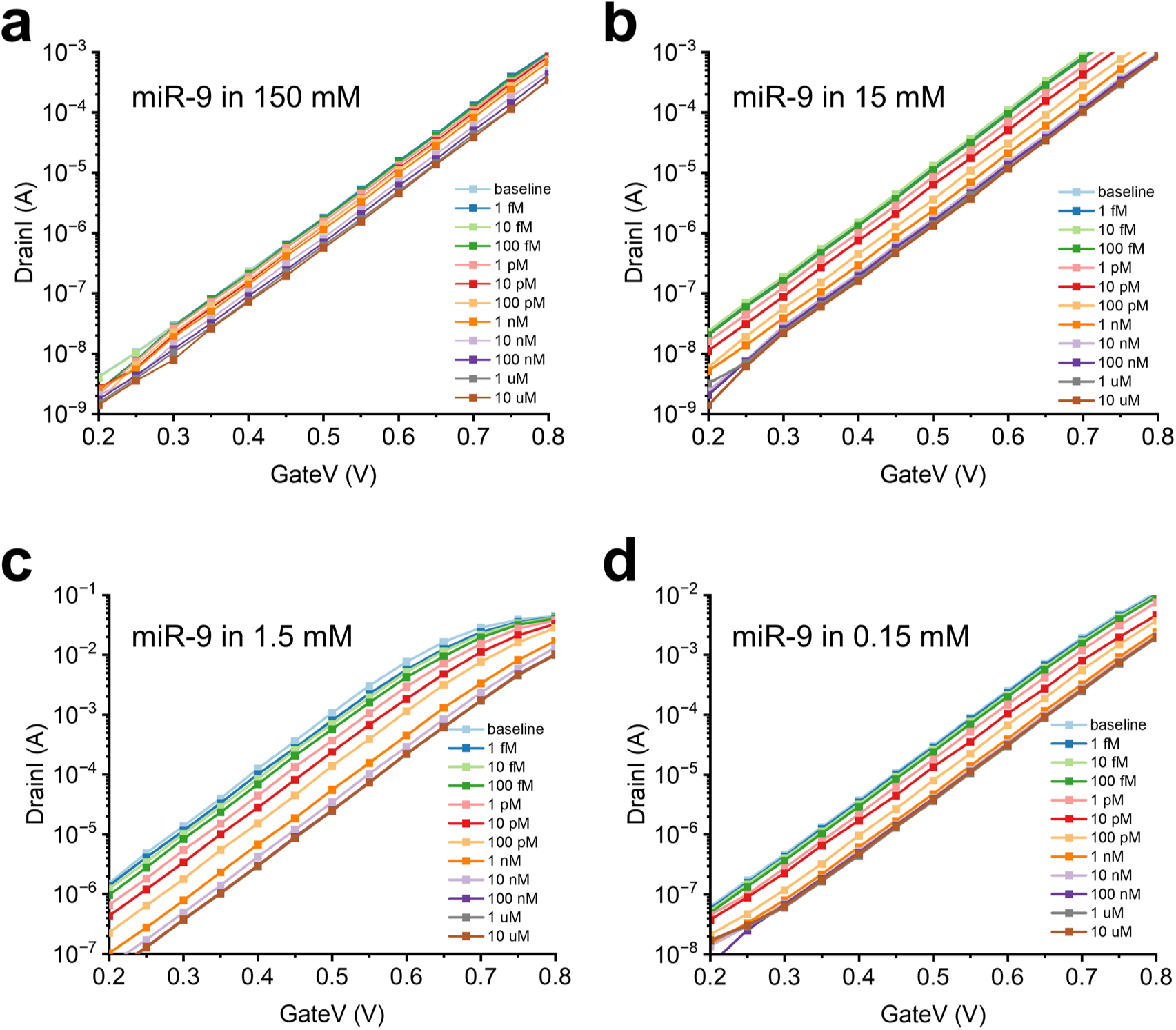
Transfer characteristics for miR-9 detection as a function of buffer ionic strength. I_DS_ vs. V_GS_ from 0.2 to 0.8 V at fixed V_DS_ = 0.1 V, recorded on probe-functionalized extended gates before incubation (baseline) and after incubation with complementary miR-9 target from 1 fM to 10 µM in (a) 150 mM, (b) 15 mM, (c) 1.5 mM, and (d) 0.15 mM ionic strength buffer.

The transfer characteristic was recorded at a fixed drain voltage V_DS_ = 0.1 V. On a semi-logarithmic scale, I_DS_ increases exponentially with V_GS_ below threshold, confirming that the device is operated in the subthreshold (weak inversion) regime of the finite-gap FET. In this regime, the drain current is governed by:

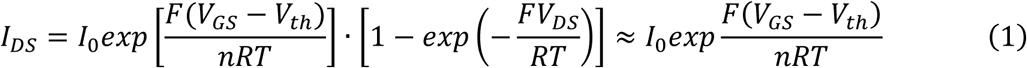

where I_0_ is the characteristic current, V_th_ the threshold voltage, RT/F ≈ 25.9 mV the thermal voltage at 300 K, and n the subthreshold ideality factor. Because V_DS_ = 0.1 V ≫ RT/F, the bracketed term approaches unity and I_DS_ depends exponentially on (V_GS_ − V_th_) alone. The subthreshold swing was extracted from the linear portion of the semi-log curve as *SS = [d(log_10_ I_DS_)/dV_GS_]^−1^ = n(RT/F) ln10*. We obtained SS ≈ 108.2 mV dec^−1^, corresponding to n ≈ 1.83, which is the value used in this report for our commercial EGFET. The elevated n beyond the ideal 59.5 mV dec^−1^ limit reflects the intrinsic capacitive non-ideality of the commercial device, the divider between its oxide and depletion capacitances together with interface trap states.^38^ It is well known that SS can be improved toward its ideal value by employing stronger electrostatic gate control through multi-gate structures, as demonstrated in FinFET and nanosheet transistors in Si CMOS.^39^ Such approaches can also be adopted here if needed. Nevertheless, a constant n is a desirable property for our platform: n is a fixed device constant and does not vary with analyte or ionic strength of the buffer, consistent with the common slope of all four panels in Figure 2.

Because ln(I_DS_) is linear in the gate potential with slope *F/(nRT)*, any binding-induced shift in the interfacial potential (the effective Zeta potential of the analyte) produces a proportional shift in the log drain current. For a negatively charged analyte on an n-channel device, the accumulated surface charge opposes the applied gate field and raises the effective threshold. Equivalently, the gating voltage is reduced by *ΔV_analyte_*, the effective Zeta potential produced by the analyte, so that *V_GS_ = V ^0^ − ΔV_analyte_*. This observation on the signal of finite-gap FET suggests the following protocol to obtain the most pertinent output signal: the applied gate voltage V_GS_^0^ is held fixed, and the baseline drain current I_B_ (before binding) and target current I_T_ (after binding) are both read at the same V_GS_ within the subthreshold region. Taking the ratio of I_B_ and I_T_ at a fixed operating point,

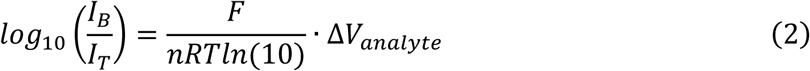

We therefore use log_10_(I_B_/I_T_), rather than the usual raw current change (I_B_ − I_T_) in earlier FET reports, as the sensing output signal: it is the device-level quantity that maps linearly onto the interfacial potential shift. Equation (2) also determines the signal cutoff for the Debye region at *ΔV_analyte_*= *RT/F*. A potential shift equal to the thermal voltage *RT/F* = 25.9 mV corresponds to log_10_(I_B_/I_T_) = 0.24 at SS ≈ 108.2 mV dec^−1^ (n = 1.83). We will demonstrate that this cutoff value of log_10_(I_B_/I_T_) is indeed the boundary between the high-noise Debye region and the low-noise GCS region and any response below this noise cutoff suffers from reproducibility issues. Figure 3 shows the response measured on this scale for all three liquid biopsy targets, on the same transducer and at the same operating point throughout. miR-9 was detected directly, through its own backbone charge. Extracellular vesicles and the protein EDN1 were detected through the silica and magnetic NP charge reporters described above. We note that, without reporters, the data for the latter two analytes fall below the GCS thermal-noise cutoff and do not provide any quantification capability--the output signal does not change with any statistical significance (more than one error bar) over 4 to 5 logs of analyte concentration. This irreproducibility is observed for all data below the cutoff for analytes both with and without reporters.

**Figure 3.**
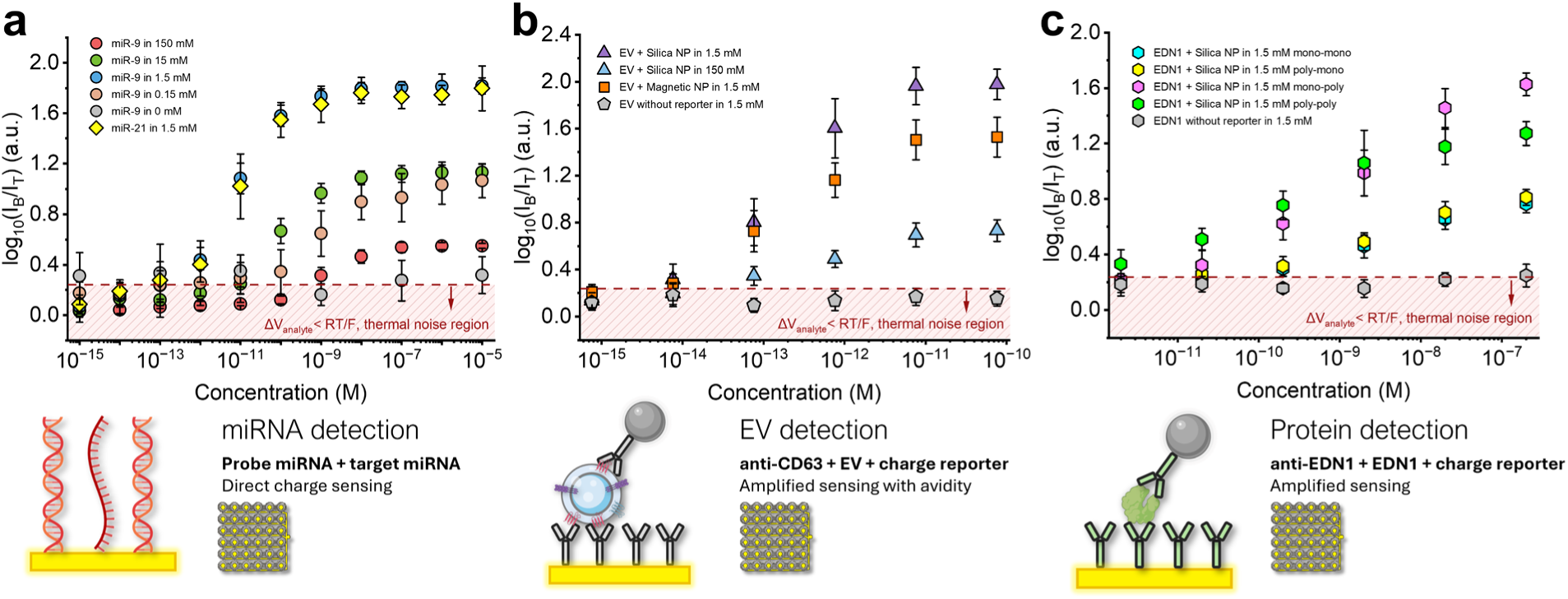
Concentration response for three classes of analyte measured on a single shared transducer. Signal log_10_(I_B_/I_T_), read at a fixed V_GS_ within the subthreshold region, versus analyte concentration. (a) Direct detection of miR-9 at buffer ionic strengths of 150, 15, 1.5, 0.15, and 0 mM, together with miR-21 at 1.5 mM. (b) EVs captured by anti-CD63 and labeled with anti-CD63 silica NP reporters at 1.5 mM and at 150 mM, with magnetic NP reporters at 1.5 mM, and without a reporter at 1.5 mM. (c) EDN1 protein at 1.5 mM in four capture-reporter antibody schemes, denoted capture-reporter: monoclonal-monoclonal, polyclonal-monoclonal, monoclonal-polyclonal and polyclonal-polyclonal, together with EDN1 incubated without a reporter. The dashed line and hatched region in each panel mark ΔV_analyte_ < RT/F, corresponding to log_10_(I_B_/I_T_) < 0.24 at n ≈ 1.83. Schematics beneath each panel show the corresponding surface chemistry and capture mode.

For data above the cutoff, the output signal increases linearly over two to three decades of analyte concentration before saturating as the probes fill. In Figure 3a, the miR-9 response is largest at 1.5 mM, while the 15 mM and 0.15 mM series are comparable. The 150 mM data barely exceed the noise cutoff, and without added salt the signal stays below log_10_(I_B_/I_T_) = 0.24 for all analyte concentrations. Taken together, these data suggest a non-monotonic sensitivity dependence on ionic strength. miR-21, measured at 1.5 mM ionic strength, provides a second miRNA sequence of similar length and charge density. Its response closely follows that of miR-9, as expected. Neither the EV nor the protein EDN1 carries enough native charge to produce sufficient signal for quantification and charge reporters are necessary in both cases. Nanoparticle tracking analysis gave 4.60 × 10^10^ particles per mL for the purified stock, equivalent to 7.64 × 10^−11^ M, and the concentration series was a 10-fold serial dilution of it (Figure S2). In Figure 3b, EVs labeled with the silica reporter gave a larger signal than those labeled with the magnetic NP at 1.5 mM, consistent with the larger zeta potential of silica. For EDN1, Figure 3c compares four capture-reporter antibody combinations at 1.5 mM. All four give the same sigmoidal form, but the reporter antibody determines the signal amplitude: a polyclonal reporter reached 1.6 and 1.3 at the highest concentration, against 0.8 for either capture antibody with a monoclonal reporter. A polyclonal reporter can bind multiple epitopes of the protein and therefore delivers more NPs to the captured proteins, whereas the capture antibody on the gate has negligible effect on the response, suggesting equal affinity (*K_D_*) for the two capture antibodies.

### GCS Theory for Noise Reduction

We employ the GCS theory to further analyze and collapse these response curves with different analytes, reporters and buffer solutions. Two links connect *ΔV_analyte_* to the surface concentration of the reporters. By treating the entire molecular layer on the electrode as a flat surface, despite the inhomogeneous charge density and the surface morphology, the Grahame equation from classical Zeta potential theory^21^ relates the interfacial potential to the effective change in surface charge density Δσ, which is the additional charge per unit area delivered to the gate due to target binding (for a symmetric univalent electrolyte),

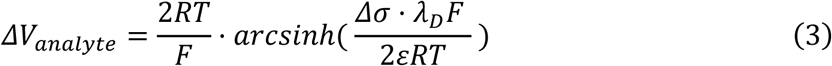

where ε is the absolute permittivity of the electrolyte, and λ_D_ the Debye length, which is set by the ionic strength of the buffer. The arcsinh form is the exact solution of the nonlinear Poisson-Boltzmann equation for the Gouy-Chapman-Stern double layer, which reduces to a linear relationship between *ΔV_analyte_* and Δσ only when the interfacial potential is small compared to *RT/F*. The surface charge density Δσ is related to the bulk analyte concentration [A] by the Langmuir isotherm^40^,

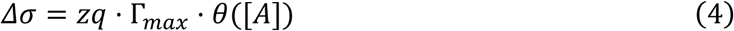

with θ([A]) = [A]/(K_D_ + [A]) being an order unity probe saturation fraction, where z is the effective number of charges carried by each bound target or reporter, q the elementary charge, Γ_max_ the surface density of binding sites at saturation, θ([A]) the fractional occupancy of those sites, [A] the target concentration, and K_D_ the equilibrium dissociation constant of the surface binding reaction.

Note that we have omitted the affinity of the reporter-analyte binding. We typically insert high concentrations of reporters so that every analyte is bound to a single reporter. In the case of EVs, we have also optimized the reporter particle size (∼50 nm) such that only one particle reporter can bind to each EV. Removal of nonspecifically bound reporters, facilitated by their large hydrodynamic size, is discussed in our earlier reports.^41^ Hence, the dissociation constant *K_D_* corresponds only to the capture of the analyte by the surface probes.

With this approximation, the effective Zeta potential becomes

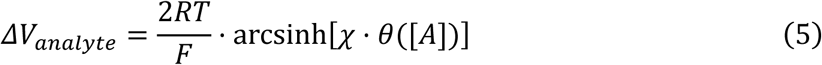

where the surface charge density due to the analyte are grouped into a characteristic length *λ_S_ ≡ 2 εRT/(zqFΓ_max_)*, often called the Gouy-Chapman or Stern length^21^. The other length scale is the usual Debey length 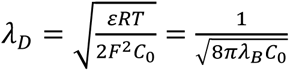 with an -1/2 scaling with respect to the bulk concentration *C*_0_ and 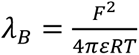 is the Bjerrum length when the electrostatic energy of two charges is equal to the thermal energy. The dimensionless gain χ ≡ λ_D_/λ_S_ hence represents the ratio between two length scales in the Debye layer, the Debye length and the Stern length for the diffusive layer. As arcsinh[*χθ*] ∼*χθ* for small *χθ*, the Debye-Hückel noise-cutoff 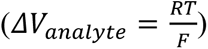 corresponds to *χθ* = ½. The Stern layer does not appear and there is only one length scale in the Debye layer for *χθ* ≪ 1. Since the fraction coverage *θ* is bound by unity, Stern layer cannot exist for χ << 1. For large χ, however, the Stern layer with thickness *λ_S_* can appear within the diffuse layer with the Debye length thickness if the fraction coverage *θ* is not too small, *χθ* ≫ 1. A large χ is hence desirable as it will reduce the critical coverage fraction to exceed the noise cutoff.

Expression (5) becomes the Donnan potential of a nanoporous membrane, after replacing the Debye length by the pore radius. As in Donnan theory for a membrane, the average mobile ion concentrations in the Debye layer (both Stern and Diffuse layer) is *C*_±_ = *C*_0_*e*^∓*F*Δ*V*^*^analyte^*^/*RT*^ for a symmetric univalent electrolyte. For small *χθ*, when the stern layer does not exist in the Debye-Hückel limit, 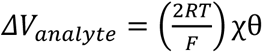 and *C*_±_= *C*_0_ (1 ± 2χθ) and the Debye layer mobile ions are sensitively dependent on the bulk concentration C_0_. In the GCS limit with large *χθ*, however, 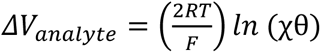 and all mobile ions in the Debye layer are counterions with a concentration of 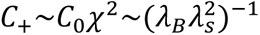 which is independent of the bulk electrolyte concentration.^29^ The mobile counterions in the Stern layer have completely compensated the dense charge on the surface and screened its high electric field. The Stern layer next to the surface is hence electrically insulated from the bulk electrolyte and hence not influenced by its variations. The significantly reduced surface field leakage minimizes counterion migration towards the surface and hence preserve the surface charge. In our earlier experiments with nanopores^30,31^, we have found that the conductance of the nanopore becomes independent of bulk ionic strength when *χθ* becomes large. These observations suggest that at the large-*χθ* GCS limit, the FET signal-to-noise ratio should be significantly higher than in the low- *χθ* Debye-Hückel limit. Some classical theories for FET charge transduction models the probe-target layer as a nanoporous membrane.^42,43^ The Debye length in expression (3) would need to be replaced by a characteristic pore radius. However, as shown by Yan et al^29^, the conclusion that the ionic concentration in the membrane becomes independent of the bulk at large Donnan potential (>>*RT/F*) remains valid independent of the pore radius, so long that it is smaller than the Debye length.

Substituting equations (3) and (4) into (2) yields an explicit relationship between the output signal and the analyte concentration [A],

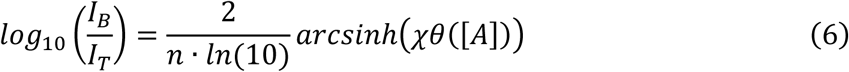

Since the subthreshold ideality factor n is fixed independently by the transfer characteristic, χ and K_D_ are the only independent parameters that relate the analyte concentration [A] to the measured signal 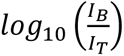. It is then clear that a large gain χ = λ_D_/λ_S_ would produce a large signal, and so would a smaller ideality factor n. At large *x*, *arcsinhx* approaches *lnx,* and we expect a linear range between the output signal and log[A] in the linear Langmuir region, if *χθ* ≫ 1 in this region. Hence, maximizing *χ* should enhance the performance of the FET sensor, both in terms of reducing noise and yielding a linear calibration. However, as the GCS limit is determined by the magnitude *χθ*, we expect both desirable features will disappear at a sufficiently low coverage *θ*. We tabulate the estimated *χ* for different probe-target and buffer ionic strengths in Table 1.

**Table 1.** Debye length λ_D_, Gouy-Chapman length λ_S_, dimensionless gain χ, and dissociation constant K_D_ for each analyte and buffer ionic strength.

| Analyte | Ionic Strength (mM) | $\lambda_D$ (nm) | $\lambda_S$ (nm) | $\chi$ | $K_D$ (M) |
| --- | --- | --- | --- | --- | --- |
| miR-9 | 0.15 | 24.82 | $5.86 \pm 0.54$ | $4.24 \pm 0.39$ | $(1.26 \pm 1.24) \times 10^{-9}$ |
| miR-9 | 1.50 | 7.85 | $0.35 \pm 0.01$ | $22.29 \pm 0.77$ | $(5.27 \pm 1.39) \times 10^{-11}$ |
| miR-9 | 15 | 2.48 | $0.48 \pm 0.02$ | $5.20 \pm 0.19$ | $(2.13 \pm 0.64) \times 10^{-10}$ |
| miR-9 | 150 | 0.78 | $0.56 \pm 0.01$ | $1.39 \pm 0.03$ | $(9.09 \pm 2.62) \times 10^{-10}$ |
| miR-21 | 1.50 | 7.85 | $0.39 \pm 0.02$ | $20.29 \pm 1.13$ | $(4.69 \pm 2.30) \times 10^{-11}$ |
| EV + silica NP | 1.50 | 7.85 | $0.23 \pm 0.02$ | $34.41 \pm 3.55$ | $(8.32 \pm 4.54) \times 10^{-13}$ |
| EV + silica NP | 150 | 0.78 | $0.37 \pm 0.04$ | $2.10 \pm 0.21$ | $(2.37 \pm 1.82) \times 10^{-13}$ |
| EV + magnetic NP | 1.50 | 7.85 | $0.59 \pm 0.08$ | $13.28 \pm 1.59$ | $(8.22 \pm 4.66) \times 10^{-13}$ |
| EDN1 protein + silica NP (mono-poly) | 1.50 | 7.85 | $0.50 \pm 0.05$ | $15.64 \pm 1.44$ | $(6.63 \pm 3.84) \times 10^{-9}$ |

For each concentration series in Figure 3, χ and K_D_ were estimated simultaneously by fitting equation (6) to the measured signal, as described in Methods. Individual fits for all measured series are shown in Figures S4–S15, with R^2^ in Table S1. Three of these are controls that show no concentration dependence, i.e., miR-9 in 0 mM, EVs without a reporter and EDN1 without a reporter. Their fitted parameters are diagnostic only and are excluded from Table 1. Table 1 lists both parameters for every analyte and buffer, together with λ_S_ obtained as λ_D_/χ. These values test the central prediction of the model. Because λ_S_ is set primarily by the target or reporter charge density, while the Debye length scales as λ_D_ ∝ I^−1/2^ for a symmetric electrolyte of ionic strength I, the gain scales as χ ∝ I^−1/2^ whereas λ_S_ is independent of ionic strength. Table 1 supports this over the upper three decades of ionic strength. For miR-9, χ = 1.39, 5.20 and 22.29 at 150, 15 and 1.5 mM, giving successive ratios of 3.74 and 4.29 per decade against the 10^1/2^ = 3.16 expected. Over these three buffers λ_S_ stayed within 0.35 to 0.56 nm, as the model requires. At 0.15 mM the trend reversed. The gain fell to 4.24, λ_S_ rose 10-fold to 5.86 nm, and K_D_ increases to (1.26 ± 1.24) × 10^−9^ M. The duplex is destabilized in this most dilute buffer, so fewer targets hybridize and Γ_max_ falls. The loss of duplex stability therefore degrades the gain as well as the affinity. Excluding this condition, λ_S_ falls between 0.23 and 0.59 nm across all analytes. Inverting λ_S_ ≡ 2εRT/(zqFΓ_max_) converts this into a saturation charge density of zΓ_max_ = 3.8 to 9.8 × 10^13^ cm^−2^, within the range of 1 to 10 × 10^12^ cm^−2^ reported for thiolated oligonucleotide monolayers on gold^44^ and well below the close-packed limit of 6 to 9 × 10^13^ cm^−2^.^45^ The dependence of the gain χ on ionic strength is therefore through λ_D_ alone, but only while the duplex remains stable.

### Universal Calibration, Detection Limit, and Dynamic Range

Equation (6) represents the universal calibration curve of the platform. Inverting it recovers the fractional occupancy directly from the measured signal,

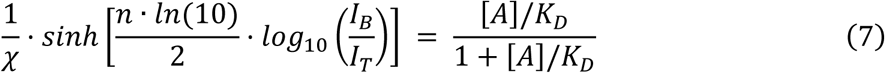

so that all the calibration curves in Figure 3, acquired for different analytes, buffers, and reporters, collapse into a single Langmuir isotherm when this χ-normalized nonlinear function of the drain current signal is plotted against the normalized concentration [A]/*K_D_* (Figure 4). This collapse into a universal calibration curve indicates that we have faithfully captured the analyte concentration-signal relationship with our GCS theory.

Plotted against log_10_([A]/K_D_), the calibration curve is a symmetric sigmoid with three segments. As the function *sinhx* is linear for small *x* but grows exponentially as *exp(x)/2* for large *x*, these different segments correspond to different dependence of the output signal on the analyte concentration. The linear and exponential scaling with respect to the output signal corresponds to the relative magnitude of *ΔV_analyte_* with respect to RT/F. Hence, the different segments of the universal curve are related to the low effective Zeta potential limit and the probe saturation limit. The low-potential Debye-Hückel limit, when *ΔV_analyte_* is smaller than RT/F and is proportional to *[A]/K_D_*, corresponds to the lower segment of this universal sigmoidal curve. The *sinhx* term can be approximated by *x* ≪ 1 in this region, corresponding to 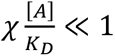. The middle linear segment in the semilog plot of the universal correlation corresponds to the GCS regime where *ΔV_analyte_*exceeds RT/F and the arcsinh crosses over to its logarithmic asymptote *ln*(2*x*). This segment requires χ[A]/K_D_ ≫ 1 while [A]/K_D_ remains small, that is K_D_/χ ≪ [A] ≪ K_D_. Saturation occurs when [A] exceeds *K_D_* and the nonlinear version of the Langmuir isotherm needs to be used. It is hence clear that the gain χ controls the transition [A] from the high-noise lower segment to the low-noise linear middle segment.

**Figure 4.**
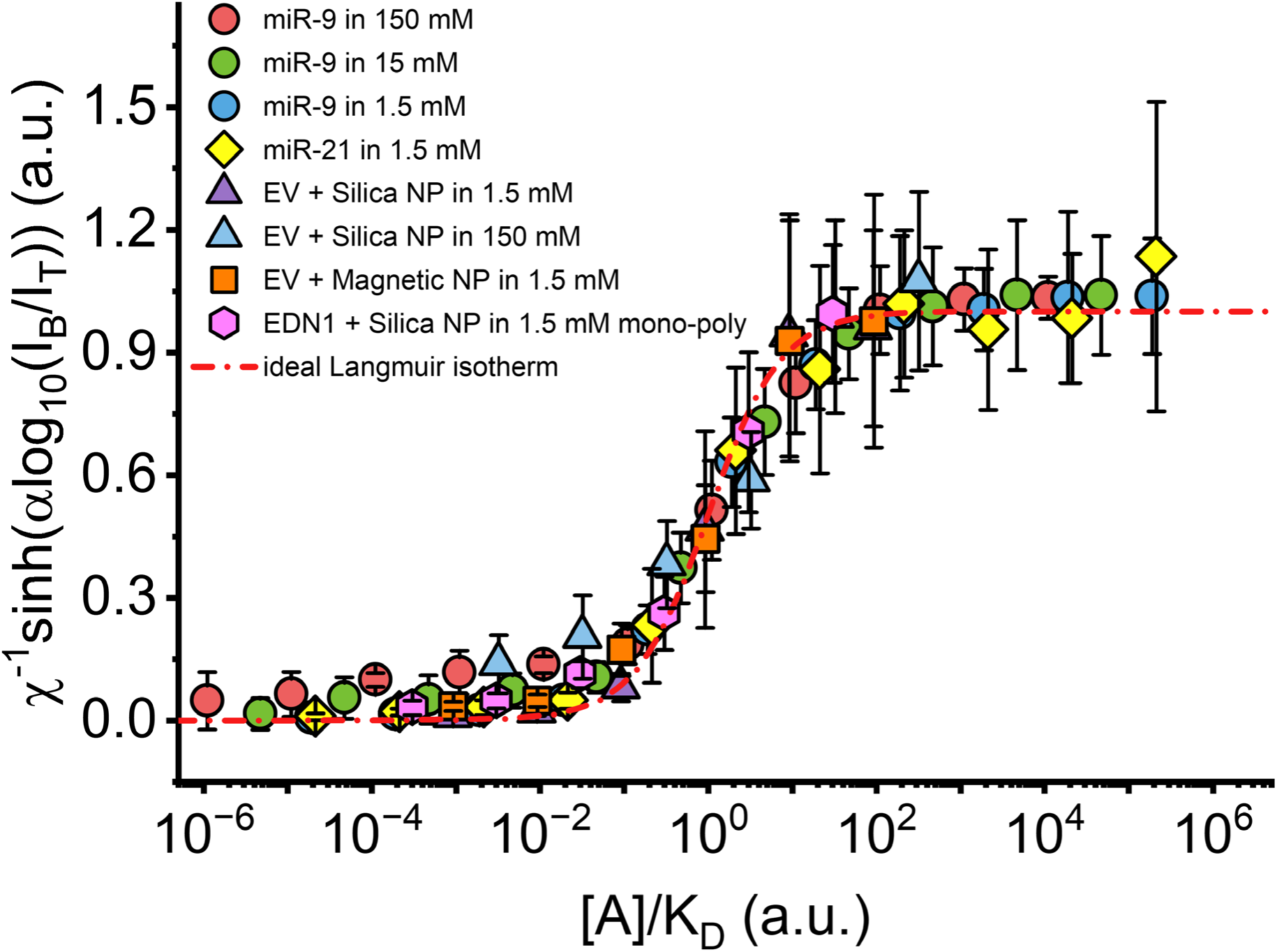
Collapse of different analytes and ionic strengths onto a single universal calibration curve. The signals of Figure 3 are inverted through the two device-level nonlinearities and plotted as χ^-1^·sinh[α·log_10_(I_B_/I_T_)], with α ≡ n·ln(10)/2 and n the subthreshold ideality factor fixed independently by the transfer characteristic, against concentration normalized by the fitted dissociation constant, [A]/K_D_. Eight analyte and condition combinations are shown. The dash-dotted line is the ideal Langmuir isotherm θ = [A]/(K_D_ + [A]).

Eight combinations of target, capture chemistry, reporter, and ionic strength, each inverted through equation (6) with only χ and *K_D_* fitted per condition, fall on the ideal Langmuir isotherm over eleven decades of normalized concentration. Before normalization, these eight datasets differ by 25-fold in gain and by four orders of magnitude in dissociation constant. They are indistinguishable after normalization and from the Langmuir isotherm. Nucleic acid, vesicle, and protein targets become indistinguishable once normalized, as do direct and reporter-mediated capture and two decades of ionic strengths. A single pair of parameters therefore converts any measurement on this platform into fractional occupancy. Slight deviation from the isotherm only occurs at the low concentration end, for miR-9 at 150 mM and the EVs labeled with the silica NP at 150 mM. These are the two lowest-gain conditions in Table 1, χ = 1.39 and 2.10, so their raw signals lie closest to the 0.24 noise cutoff. Both are measured at physiological ionic strength, where screening is strongest.

Taking the thermal voltage RT/F as the noise cutoff, the detection limit is the concentration at which the recovered signal emerges from the lower branch, χ·θ ∼ 1, and at low coverage θ ≈ [A]/*K_D_*, so

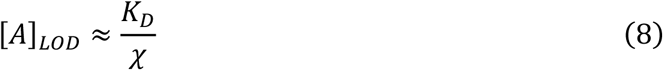

and the dynamic range of the linear middle segment ranges from this limit up to saturation, spanning orders of log_10_χ + 1 decades (roughly 2 to 3 logs for our optimized χ). Both LOD and dynamic range improve as χ increases, and the signal-to-noise ratio grows logarithmically with [A]/K_D_ through the linear region.

These explicit predictions of detection limit and dynamic range allow us to optimize the sensor platform. Lowering the ionic strength enlarges λ_D_ and hence χ, although very low ionic strength can also change the binding affinity *K_D_* through the loss of bound target. So an optimum ionic strength may exist for a given probe-target pair. Multivalent capture, as when a vesicle binds through several surface epitopes, lowers *K_D_*. A highly charged reporter raises the surface charge density and lowers λ_S_: the larger fitted gain for the silica NP reporter (χ ≈ 34) relative to the magnetic NP (χ ≈ 13) reflects exactly this, consistent with their measured zeta potentials. Probe density acts on both parameters at once: a denser probe layer raises Γ_max_ and so lowers λ_S_, while it also lowers K_D_ wherever a single target can engage several probes in a multi-valent capture mechanism, so probe density should be pushed as high as the surface chemistry allows.

Data in Figure 5 verify our theory on the low-noise GCS limit, when *χθ* becomes large, and quantify our LOD optimization effort. In Figure 5a the signal-to-noise ratio for miR-9 stays within the noise band until [A]χ/K_D_ approaches unity and rises thereafter, once the data depart from the lower Debye-Hückel branch with *χθ* ≪ 1. The 1.5 mM series reach the highest values, the 0.15 mM series barely exceed the threshold, having neither the largest gain nor the tightest binding, and without added salt the signal never crosses S/N = 3.3. A large χ alone therefore does not guarantee a better measurement, because the gain is only useful while enough target stays bound to keep χθ above unity. Figure 5b shows the signal behavior for vesicles: EVs labeled with the silica NP reach S/N ≈ 15 and those labeled with the magnetic NP ≈ 9, in the order of their gains, while the same EVs reach only ≈ 8 at 150 mM and unlabeled EVs remain in the noise band at every concentration. Figure 5c corresponds to the optimization for EDN1. The mono-poly scheme reaches S/N ≈ 20, poly-poly and poly-mono are indistinguishable at ≈ 15, and mono-mono ≈ 12. Only the mono-poly scheme separates clearly from the rest, as it does in the amplitudes of Figure 3c. Figure 5d resolves the competition between binding affinity and signal amplification for miR-9 detection. At low ionic strengths, hybridization affinity is expected to deteriorate due to electrostatic repulsion between the like-charged analyte and oligo probe, but the output signal is expected to increase due to decreasing Debye screening. Accordingly, the measured gain parameter χ increases as the ionic strength drops from 150 to 1.5 mM, following λ_D_, but it decreases again at 0.15 mM, and *K_D_* and the detection limit both reach their minimum at the same 1.5 mM optimum. Above 1.5 mM, Debye screening reduces the amplitude of signal transduction. Below it, the duplex is destabilized, so both the affinity and the gain degrade. The detection limit is optimized at an intermediate ionic strength, 1.5 mM for this probe–target pair, where equation (8) gives K_D_/χ = 2.4 × 10^−12^ M against a measured 1 × 10^−12^ M, the lowest concentration reaching S/N ≥ 3.3, so the predicted and measured limits agree to within a factor of 2.4.

**Figure 5.**
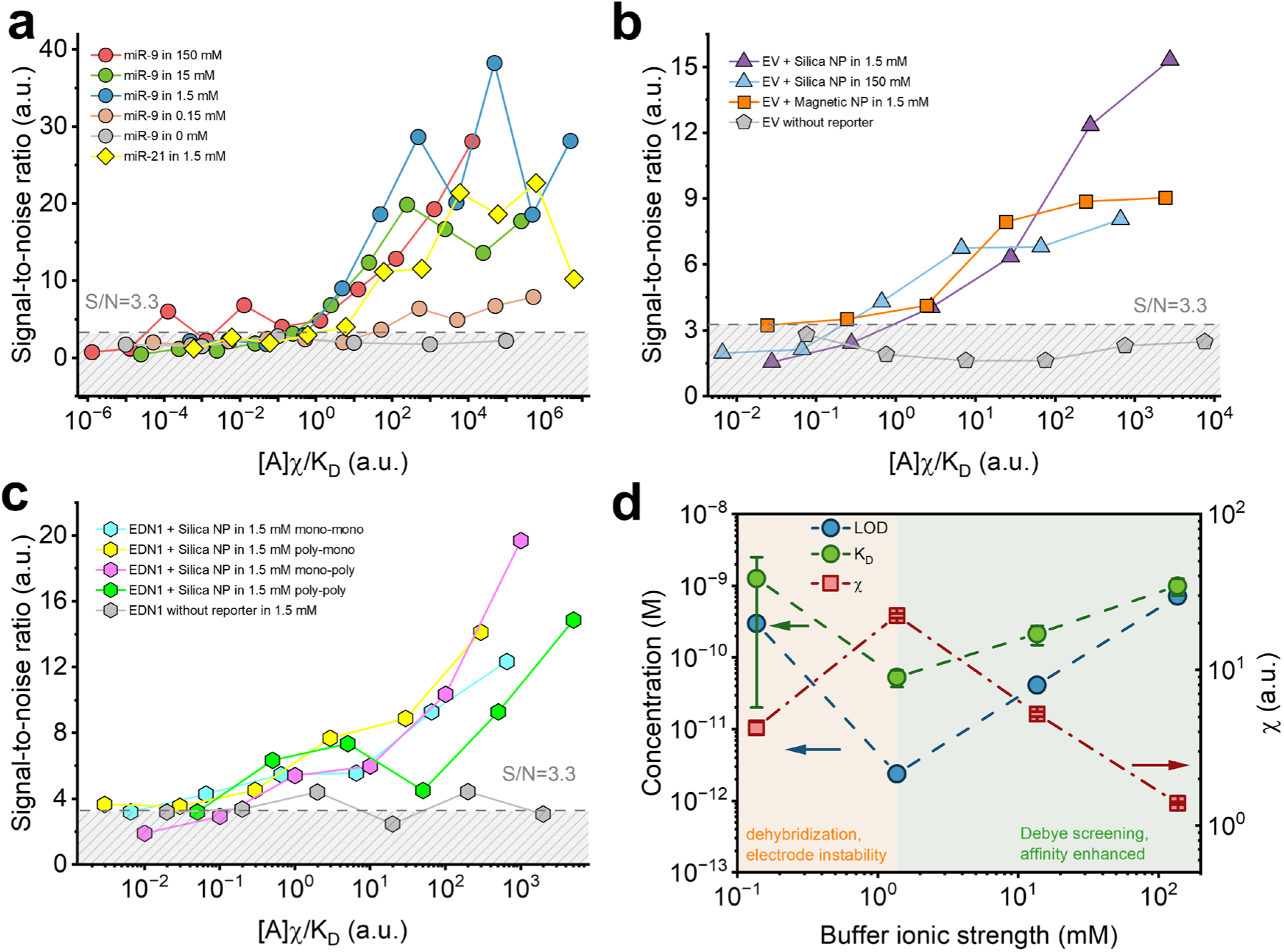
Signal-to-noise ratio and the optimum ionic strength. Signal-to-noise ratio versus reduced concentration [A]χ/K_D_ for (a) miR-9 at 150, 15, 1.5, 0.15, and 0 mM and miR-21 at 1.5 mM, (b) EVs with the silica NP at 1.5 and 150 mM, with the magnetic NP at 1.5 mM, and without a reporter, and (c) EDN1 at 1.5 mM in the four capture-reporter schemes and without a reporter. Dashed lines mark S/N = 3.3, and hatched bands mark the region indistinguishable from noise. (d) For direct miR-9 detection, predicted limit of detection K_D_/χ and dissociation constant K_D_ (left axis) and gain χ (right axis) versus buffer ionic strength. Shaded regions mark dehybridization and electrode instability below 1.5 mM and Debye screening with enhanced affinity above it.

### Specificity, Probe Density, and Multi-Valent Avidity

Figure 6a examines the specificity of miRNA detection for the EGFET. The miR-9 probe reports an output of 0.44 with miR-9 and a significantly lower signal of 0.05 with miR-21, while the miR-21 probe yields 0.44 with its miR-21 target and 0.07 with miR-9. All cross-reactive signals fall below the noise cutoff. The two matched pairs give identical signals, as expected for sequences of equal length and charge. Mismatches can also be discerned by the EGFET. The target presented to the miR-9 probe varied from a perfect complement to targets carrying 1, 2, and 3 mismatched bases (Figure 6b). The perfect target gave 0.34 and a single mismatch 0.31, both above the noise cutoff, whereas two and three mismatches gave 0.16 and 0.09, both below it (Figure 6c). Discrimination of 2 or more mismatches is hence expected to be highly reproducible.

**Figure 6.**
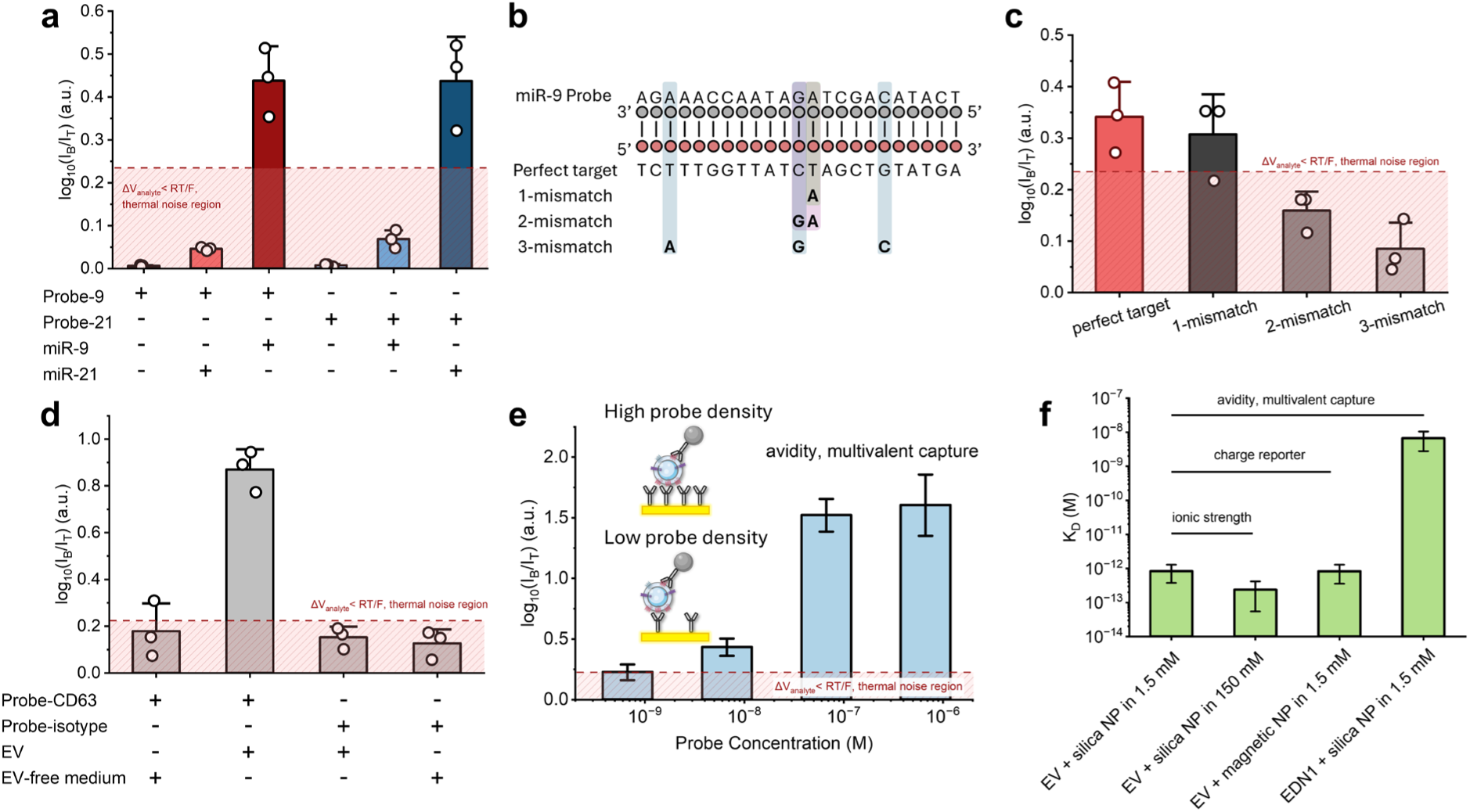
Specificity, probe density and avidity. (a) Cross-reactivity panel: gates functionalized with the miR-9 or the miR-21 probe, each exposed to miR-9 (1 × 10^−12^ M), to miR-21 (1 × 10^−12^ M), and to no target. (b) Sequences of the miR-9 probe and of the perfect, 1-, 2-, and 3-mismatch targets, with mismatched bases highlighted. (c) Response of miR-9 probe gates to a perfectly complementary target and to targets with 1-, 2-, and 3-base mismatches, all at 1 × 10^−12^ M. (d) Control panel comparing anti-CD63 and isotype-control gates against EV-containing medium at 7.64 × 10^−14^ M and EV-free medium, each labeled with the silica NP reporter. (e) Signal for anti-CD63-captured EVs at 7.64 × 10^−13^ M as a function of the anti-CD63 concentration used to functionalize the gate: 6.7 × 10^−7^ M and its 1:10, 1:100, and 1:1000 dilutions, with the silica NP reporter throughout. Insets illustrate high and low probe density. (f) Dissociation constant K_D_ for EVs with silica reporters at 1.5 mM and at 150 mM, EVs with magnetic reporters at 1.5 mM, and EDN1 protein with silica reporters at 1.5 mM. Horizontal bars mark the pairwise comparisons that isolate ionic strength, charge reporter, and avidity.

The same test for vesicles is in Figure 6d. Anti-CD63 gates exposed to EVs at 7.64 × 10^−14^ M gave 0.87, whereas the isotype-control gates and the EV-free medium all remained within the noise cutoff at 0.24. Capture is therefore antibody-mediated and not the result of nonspecific adsorption of vesicles or of the charge reporter.

Figure 6e isolates probe density. EVs at 7.64 × 10^−13^ M were measured on sensor electrodes functionalized with a 6.7 × 10^−7^ M anti-CD63 stock and its 1:10, 1:100, and 1:1000 dilutions, each resulting in a different probe density. The signal rose from 0.22 at the 1:1000 dilution to 1.61 at the undiluted stock, with most of the increase between the 1:100 and 1:10 dilutions, that is, between 6.7 × 10^−9^ and 6.7 × 10^−8^ M, and a plateau thereafter. A dense antibody layer allows a single vesicle to engage several probes, thus allowing multivalent capture. This avidity lowers *K_D_* such that it falls below 7.64 × 10^−13^ M at high probe density and above it at low probe density.

We compared the three key factors affecting the sensitivity of EV detection (Figure 6f). Changing the ionic strength reduces *K_D_* by a factor of 3.5, from (8.32 ± 4.54) × 10^−13^ M at an ionic strength of 1.5 mM to (2.37 ± 1.82) × 10^−13^ M at 150 mM. Debye screening improves binding affinity even as it reduces gain, which is the trade-off also observed for miRNA detection in Figure 5d. However, this correction is much smaller for EVs, reflecting their weaker charge. Changing the reporter from silica to magnetic beads with a smaller charge leaves KD unchanged at (8.22 ± 4.66) × 10^−13^ M against the silica reporter at the same ionic strength, so the reporter acts principally on χ rather than on binding. In contrast, changing the target from a vesicle to a protein in the monoclonal-capture/polyclonal-reporter scheme moves it by a factor of about 8,000, to (6.63 ± 3.84) × 10^−9^ M, because a protein presents a single epitope and cannot bind multivalently. Avidity is thus the dominant determinant of *K_D_* for EV detection, with ionic strength having a distant secondary effect. Multivalent capture is hence why EVs reach femtomolar detection limits with a gain only twice that of the protein assay.

## CONCLUSIONS

We developed a 36-well extended-gate FET biosensor array in which gold electrodes patterned on a polycarbonate substrate are connected to the gate of a single shared commercial MOSFET. The array carries the surface chemistry while the transducer is reused indefinitely, so every measurement reported here was made on the same transistor at the same operating point. A finite-bandgap channel removes the electronic noise that limits zero-bandgap devices, leaving electrolyte fluctuation as the only noise source at the interface. We have demonstrated that this fluctuation only affects the gating voltage in the linear Debye-Hückel limit. Once the interfacial potential exceeds the thermal voltage RT/F, a Stern layer appears to insulate the surface from the electrolyte fluctuations. The measured Gouy-Chapman-Stern output is highly reproducible for the three liquid biopsy analytes we tested. The GCS theory produces two parameters, the gain χ and the dissociation constant KD, whose determination allows us to collapse all calibration curves into a universal sigmoid corresponding to the Langmuir isotherm. Due to the different Zeta potential scaling with respect to the charge in the Debye-Hückel and GCS regimes, the noisy Debye regime is conveniently related to the lower segment of the sigmoid with a very specific cutoff on the output signal defined by our theory. The theory also offers an explicit prediction of the LOD and the linear dynamic range based on the two parameters. These predictions allow for optimization of the buffer ionic strength, reporter, and critical probe density for each target. We believe our results will finally enable wide use of the finite-bandgap EGFET sensor, particularly in a multi-array format that maximizes its advantages. A clinical liquid biopsy blood test with multiple analyte targets and multiple samples would be one such application.

## Author Information

### Authors

Feng Gao – Department of Chemical and Biomolecular Engineering, University of Notre Dame, Indiana, 46556, USA

Tiger Haoran Shi – Department of Chemical and Biomolecular Engineering, University of Notre Dame, Indiana, 46556, USA

Tyler Moorman – Department of Chemical and Biomolecular Engineering, University of Notre Dame, Indiana, 46556, USA

Sizhe Ma – Department of Electrical Engineering, University of Notre Dame, Indiana, 46556, USA

Kai Ni – Department of Electrical Engineering, University of Notre Dame, Indiana, 46556, USA

Satyajyoti Senapati – Department of Chemical and Biomolecular Engineering, University of Notre Dame, Indiana, 46556, USA; https://orcid.org/0000-0001-7999-1561

## Acknowledgement

H.C.C. and S.S. acknowledge the support from the NIH, 1R21AI180713-01A1. We thank the laboratory of Prof. Robert J. Coffey at the Department of Medicine (Gastroenterology and Cell & Developmental Biology), Vanderbilt University Medical Center, for providing DiFi conditioned medium. We thank the Materials Characterization Facility (MCF), supported by Notre Dame Research, for use of the Zetasizer, and the Harper Cancer Research Institute, University of Notre Dame, for use of the NanoSight NS300

## References

(1) Janićijević, Ž.; Nguyen-Le, T.-A.; Baraban, L. Extended-Gate Field-Effect Transistor Chemo- and Biosensors: State of the Art and Perspectives. Nanotechnol. 2023, 3–4, 100025.

(2) Janićijević, Ž.; Baraban, L. Integration Strategies and Formats in Field-Effect Transistor Chemo- and Biosensors: A Critical Review. ACS Sens. 2025, 10 (4), 2431–2452.

(3) Ono, T.; Okuda, S.; Ushiba, S.; Kanai, Y.; Matsumoto, K. Challenges for Field-Effect-Transistor-Based Graphene Biosensors. Materials 2024, 17 (2), 333.

(4) Kumar, S.; Sinclair, J. A.; Shi, T.; Kim, G.; Zhu, R.; Gasper, G.; Wang, Y.; Higginbotham, J. N.; Zhang, Q.; Jeppesen, D. K.; Tutanov, O.; Hamilton, M.; Franklin, J. L.; Charest, A.; Coffey, R. J.; Senapati, S.; Chang, H.-C. Surface Markers on Supermeres Outperform Extracellular Vesicles in Colorectal Cancer Diagnosis. Sci. Rep. 2026, 16 (1), 5989.

(5) Bergveld, P. Development of an Ion-Sensitive Solid-State Device for Neurophysiological Measurements. IEEE Trans. Biomed. Eng. 1970, *BME-17* (1), 70–71.

(6) Cui, Y.; Wei, Q.; Park, H.; Lieber, C. M. Nanowire Nanosensors for Highly Sensitive and Selective Detection of Biological and Chemical Species. Science 2001, 293 (5533), 1289–1292.

(7) Hwang, M. T.; Heiranian, M.; Kim, Y.; You, S.; Leem, J.; Taqieddin, A.; Faramarzi, V.; Jing, Y.; Park, I.; Van Der Zande, A. M.; Nam, S.; Aluru, N. R.; Bashir, R. Ultrasensitive Detection of Nucleic Acids Using Deformed Graphene Channel Field Effect Biosensors. Nat. Commun. 2020, 11 (1), 1543.

(8) Béraud, A.; Sauvage, M.; Bazán, C. M.; Tie, M.; Bencherif, A.; Bouilly, D. Graphene Field-Effect Transistors as Bioanalytical Sensors: Design, Operation and Performance. The Analyst 2021, 146 (2), 403–428.

(9) Li, Y.; Peng, Z.; Holl, N. J.; Hassan, Md. R.; Pappas, J. M.; Wei, C.; Izadi, O. H.; Wang, Y.; Dong, X.; Wang, C.; Huang, Y.-W.; Kim, D.; Wu, C. MXene–Graphene Field-Effect Transistor Sensing of Influenza Virus and SARS-CoV-2. ACS Omega 2021, 6 (10), 6643–6653.

(10) Alvandi, H.; Asadi, F.; Rezayan, A. H.; Hajghassem, H.; Rahimi, F. Ultrasensitive Biosensor Based on MXene-GO Field-Effect Transistor for the Rapid Detection of Endotoxin and Whole-Cell E. Coli in Human Blood Serum. Anal. Chim. Acta 2025, 1348, 343816.

(11) Nisar, S.; Dastgeer, G.; Shazad, Z. M.; Zulfiqar, M. W.; Rasheed, A.; Iqbal, M. Z.; Hussain, K.; Rabani, I.; Kim, D.; Irfan, A.; Chaudhry, A. R. 2D Materials in Advanced Electronic Biosensors for Point-of-Care Devices. Adv. Sci. 2024, 11 (31), 2401386.

(12) Genco, E.; Modena, F.; Sarcina, L.; Björkström, K.; Brunetti, C.; Caironi, M.; Caputo, M.; Demartis, V. M.; Di Franco, C.; Frusconi, G.; Haeberle, L.; Larizza, P.; Mancini, M. T.; Österbacka, R.; Reeves, W.; Scamarcio, G.; Scandurra, C.; Wheeler, M.; Cantatore, E.; Esposito, I.; Macchia, E.; Torricelli, F.; Viola, F. A.; Torsi, L. A Single-Molecule Bioelectronic Portable Array for Early Diagnosis of Pancreatic Cancer Precursors. Adv. Mater. 2023, 35 (42), 2304102.

(13) Hung, S.-C.; Hung, K.-C.; Lin, C. K. Extended-Gate FET Biosensor Utilizing Ionic Strength Engineering for Sensitive Prostate-Specific Antigen Detection. Talanta 2026, 304, 129531.

(14) Pullano, S. A.; Critello, C. D.; Mahbub, I.; Tasneem, N. T.; Shamsir, S.; Islam, S. K.; Greco, M.; Fiorillo, A. S. EGFET-Based Sensors for Bioanalytical Applications: A Review. Sensors 2018, 18 (11), 4042.

(15) Fu, W.; Feng, L.; Panaitov, G.; Kireev, D.; Mayer, D.; Offenhäusser, A.; Krause, H.-J. Biosensing near the Neutrality Point of Graphene. Sci. Adv. 2017, 3 (10), e1701247.

(16) Loan, P. T. K.; Wu, D.; Ye, C.; Li, X.; Tra, V. T.; Wei, Q.; Fu, L.; Yu, A.; Li, L.-J.; Lin, C.-T. Hall Effect Biosensors with Ultraclean Graphene Film for Improved Sensitivity of Label-Free DNA Detection. Biosens. Bioelectron. 2018, 99, 85–91.

(17) Senapati, S.; Slouka, Z.; Shah, S. S.; Behura, S. K.; Shi, Z.; Stack, M. S.; Severson, D. W.; Chang, H.-C. An Ion-Exchange Nanomembrane Sensor for Detection of Nucleic Acids Using a Surface Charge Inversion Phenomenon. Biosens. Bioelectron. 2014, 60, 92–100.

(18) Slouka, Z.; Senapati, S.; Chang, H.-C. Microfluidic Systems with Ion-Selective Membranes. Annu. Rev. Anal. Chem. 2014, 7 (1), 317–335.

(19) Gao, X. P. A.; Zheng, G.; Lieber, C. M. Subthreshold Regime Has the Optimal Sensitivity for Nanowire FET Biosensors. Nano Lett. 2010, 10 (2), 547–552.

(20) Kesler, V.; Murmann, B.; Soh, H. T. Going beyond the Debye Length: Overcoming Charge Screening Limitations in Next-Generation Bioelectronic Sensors. ACS Nano 2020, 14 (12), 16194–16201.

(21) Chang, H.-C.; Yeo, L. Y. Electrokinetically Driven Microfluidics and Nanofluidics. Camb. Univ. Press. 2010.

(22) Stern, E.; Wagner, R.; Sigworth, F. J.; Breaker, R.; Fahmy, T. M.; Reed, M. A. Importance of the Debye Screening Length on Nanowire Field Effect Transistor Sensors. Nano Lett. 2007, 7 (11), 3405–3409.

(23) Swenson, C. S.; Lackey, H. H.; Reece, E. J.; Harris, J. M.; Heemstra, J. M.; Peterson, E. M. Evaluating the Effect of Ionic Strength on PNA:DNA Duplex Formation Kinetics. RSC Chem. Biol. 2021, 2 (4), 1249–1256.

(24) Susa, F.; Arnaboldi, L.; De Giorgis, V.; Ghirimoldi, M.; Barberis, E.; Brucale, M.; Valle, F.; Malerba, M.; Vallino, M.; Musicò, A.; Frigerio, R.; Gori, A.; Mugoni, V.; Voena, C.; Pisano, R.; Pirri, F.; Arpicco, S.; Manfredi, M.; Limongi, T. Osmotic Remodeling of Extracellular Vesicles for Precision Nanomedicine. Small 2026, e10793.

(25) Manning, G. S. Limiting Laws and Counterion Condensation in Polyelectrolyte Solutions I. Colligative Properties. J. Chem. Phys. 1969, 51 (3), 924–933.

(26) Plouraboué, F.; Chang, H.-C. Symmetry Breaking and Electrostatic Attraction between Two Identical Surfaces. Phys. Rev. E 2009, 79 (4), 041404.

(27) Macchia, E.; Torricelli, F.; Caputo, M.; Sarcina, L.; Scandurra, C.; Bollella, P.; Catacchio, M.; Piscitelli, M.; Di Franco, C.; Scamarcio, G.; Torsi, L. Point-Of-Care Ultra-Portable Single-Molecule Bioassays for One-Health. Adv. Mater. 2024, 36 (13), 2309705.

(28) Moreira, A. G.; Netz, R. R. Simulations of Counterions at Charged Plates. Eur. Phys. J. E 2002, 8 (1), 33–58.

(29) Yan, Y.; Wang, L.; Xue, J.; Chang, H.-C. Ion Current Rectification Inversion in Conic Nanopores: Nonequilibrium Ion Transport Biased by Ion Selectivity and Spatial Asymmetry. J. Chem. Phys. 2013, 138 (4), 044706.

(30) Yossifon, G.; Mushenheim, P.; Chang, Y.-C.; Chang, H.-C. Eliminating the Limiting-Current Phenomenon by Geometric Field Focusing into Nanopores and Nanoslots. Phys. Rev. E 2010, 81 (4), 046301.

(31) Chang, H.-C.; Yossifon, G.; Demekhin, E. A. Nanoscale Electrokinetics and Microvortices: How Microhydrodynamics Affects Nanofluidic Ion Flux. Annu. Rev. Fluid Mech. 2012, 44 (1), 401–426.

(32) Sensale, S.; Ramshani, Z.; Senapati, S.; Chang, H.-C. Universal Features of Non-Equilibrium Ionic Currents through Perm-Selective Membranes: Gating by Charged Nanoparticles/Macromolecules for Robust Biosensing Applications. J. Phys. Chem. B 2021, 125 (7), 1906–1915.

(33) Kumar, S.; Maniya, N.; Wang, C.; Senapati, S.; Chang, H.-C. Quantifying PON1 on HDL with Nanoparticle-Gated Electrokinetic Membrane Sensor for Accurate Cardiovascular Risk Assessment. Nat. Commun. 2023, 14 (1), 557.

(34) Vauquelin, G.; Charlton, S. J. Exploring Avidity: Understanding the Potential Gains in Functional Affinity and Target Residence Time of Bivalent and Heterobivalent Ligands. Br. J. Pharmacol. 2013, 168 (8), 1771–1785.

(35) Oostindie, S. C.; Lazar, G. A.; Schuurman, J.; Parren, P. W. H. I. Avidity in Antibody Effector Functions and Biotherapeutic Drug Design. Nat. Rev. Drug Discov. 2022, 21 (10), 715–735.

(36) Landry, J. P.; Ke, Y.; Yu, G.-L.; Zhu, X. D. Measuring Affinity Constants of 1450 Monoclonal Antibodies to Peptide Targets with a Microarray-Based Label-Free Assay Platform. J. Immunol. Methods 2015, 417, 86–96.

(37) Ko, S. Y.; Lee, W.; Naora, H. Harnessing microRNA-Enriched Extracellular Vesicles for Liquid Biopsy. Front. Mol. Biosci. 2024, 11, 1356780.

(38) Barboni, L.; Siniscalchi, M.; Sensale-Rodriguez, B. TFET-Based Circuit Design Using the Transconductance Generation Efficiency $ {g}_{m}/ {I}_{d}$ Method. IEEE J. Electron Devices Soc. 2015, 3 (3), 208–216.

(39) Yoon, J.-S.; Baek, R.-H. Device Design Guideline of 5-Nm-Node FinFETs and Nanosheet FETs for Analog/RF Applications. IEEE Access 2020, 8, 189395–189403.

(40) Langmuir, I. THE ADSORPTION OF GASES ON PLANE SURFACES OF GLASS, MICA AND PLATINUM. J. Am. Chem. Soc. 1918, 40 (9), 1361–1403.

(41) Kumar, S.; Senapati, S.; Chang, H.-C. Extracellular Vesicle and Lipoprotein Diagnostics (ExoLP-Dx) with Membrane Sensor: A Robust Microfluidic Platform to Overcome Heterogeneity. Biomicrofluidics 2024, 18 (4), 041301.

(42) Schasfoort, R. B. M.; Bergveld, P.; Kooyman, R. P. H.; Greve, J. Possibilities and Limitations of Direct Detection of Protein Charges by Means of an Immunological Field-Effect Transistor. Anal. Chim. Acta 1990, 238, 323–329.

(43) Kaisti, M. Detection Principles of Biological and Chemical FET Sensors. Biosens. Bioelectron. 2017, 98, 437–448.

(44) Steel, A. B.; Herne, T. M.; Tarlov, M. J. Electrochemical Quantitation of DNA Immobilized on Gold. Anal. Chem. 1998, 70 (22), 4670–4677.

(45) Steel, A. B.; Levicky, R. L.; Herne, T. M.; Tarlov, M. J. Immobilization of Nucleic Acids at Solid Surfaces: Effect of Oligonucleotide Length on Layer Assembly. Biophys. J. 2000, 79 (2), 975–981.

