## Supplementary Information for "Low-Noise Finite-Bandgap FET Biosensors in the Nonlinear Gouy-Chapman-Stern Region: A Universal Calibration Curve"

**Materials and Reagents**

1-ethyl-3-(3-dimethylaminopropyl) carbodiimide (EDC, PI22980), Sulfo N-hydroxysulfosuccinimide (NHS, PI24510), and 0.1 M 2-(N-morpholino) ethanesulfonic acid (MES) buffer pack (J62231.AP) were purchased from Thermo Fisher Scientific (Waltham, MA, USA). 11-Mercaptoundecanoic acid (11-MUA, 450561-5G), 200 proof ethanol (E7023-500ML), and 1× DPBS were purchased from Sigma Aldrich (St. Louis, MO, USA). Synthetic probes of miR-21 and miR-9, and their fully matched complementary sequences and sequences with one, two, and three base mismatches at different locations, were purchased from IDT (Coralville, IA, USA). CD63 monoclonal antibody (mouse IgG1, 67605-1-Ig) and the isotype-matched mouse IgG1 control antibody (66360-1-Ig) were purchased from Proteintech (Rosemont, IL, USA). Recombinant EDN1 protein (H00001906-P01) and EDN1 monoclonal antibody (H00001906-M01) were purchased from Norvus Biologicals (Colorado, USA). Rabbit polyclonal antibody to endothelin-1 (ab117757) was purchased from Abcam (Boston, USA). Carboxyl-terminated silica NPs (53.1 nm, C-SIO-0.05COOH, 140510-10) were purchased from microspheres-nanospheres.com (Cold Spring, NY, USA). Carboxyl-terminated superparamagnetic NPs (200 nm, SuperMag Carboxyl Beads, SC0200-002) were purchased from Ocean NanoTech (San Diego, CA, USA). An n-type enhancement-mode metal-oxide-semiconductor field-effect transistor (G30N02T, Wuxi Goford Semiconductor) was purchased from DigiKey (Thief River Falls, MN, USA). Conditioned medium from DiFi human colorectal carcinoma cells, the source of the extracellular vesicles used in this study, was provided by Dr. Robert J. Coffey (Vanderbilt University, Nashville, TN, USA). Epoxy resin (Quik-Cast) and polycarbonate sheets were purchased from TAP Plastics (San Leandro, CA, USA).

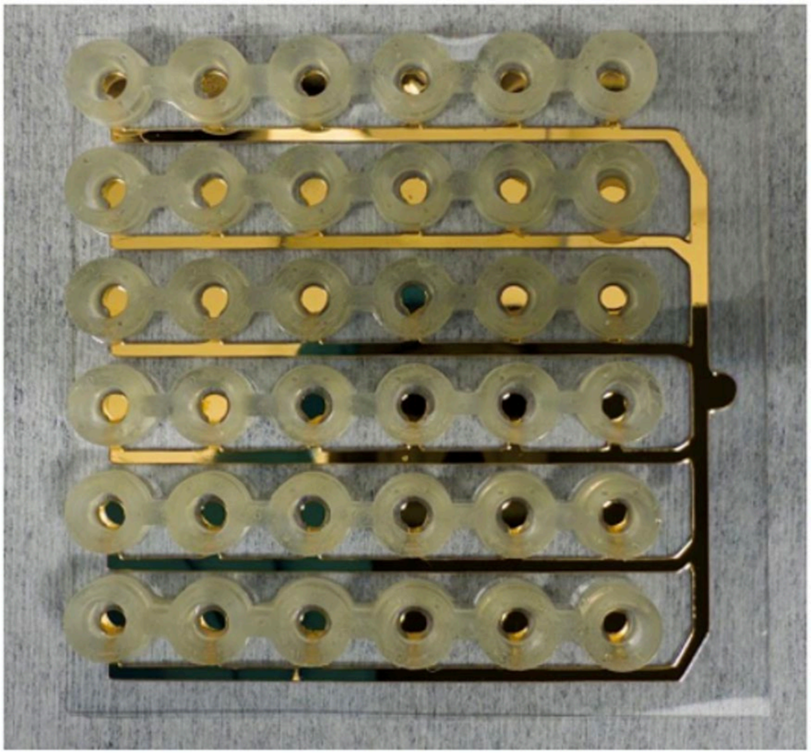

Figure S1. Photograph of the fabricated 36-well extended-gate array. The six-by-six array contains gold sensing pads surrounded by individual wells. Patterned gold traces connect the sensing pads to the common extended-gate contact.

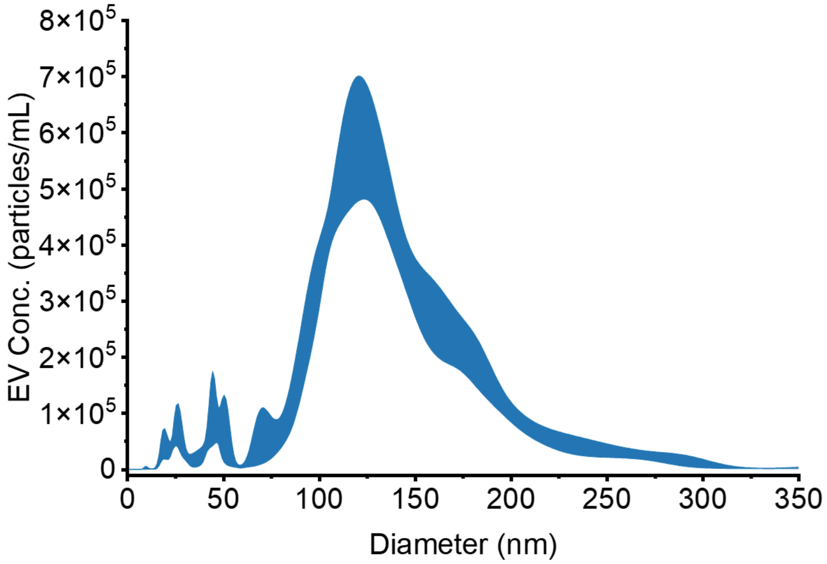


Figure S2. Size distribution of purified extracellular vesicles. Nanoparticle tracking analysis of extracellular vesicles isolated from DiFi conditioned medium. Particle concentration is plotted as a function of hydrodynamic diameter. The measurement was made on a 1000-fold dilution of the stock preparation and gave $4.60 \times 10^{7}$ particles per mL, so the undiluted stock contained $4.60 \times 10^{10}$ particles per mL, equivalent to $7.64 \times 10^{-11}$ M. All EV concentrations reported in this work are serial tenfold dilutions of this stock.

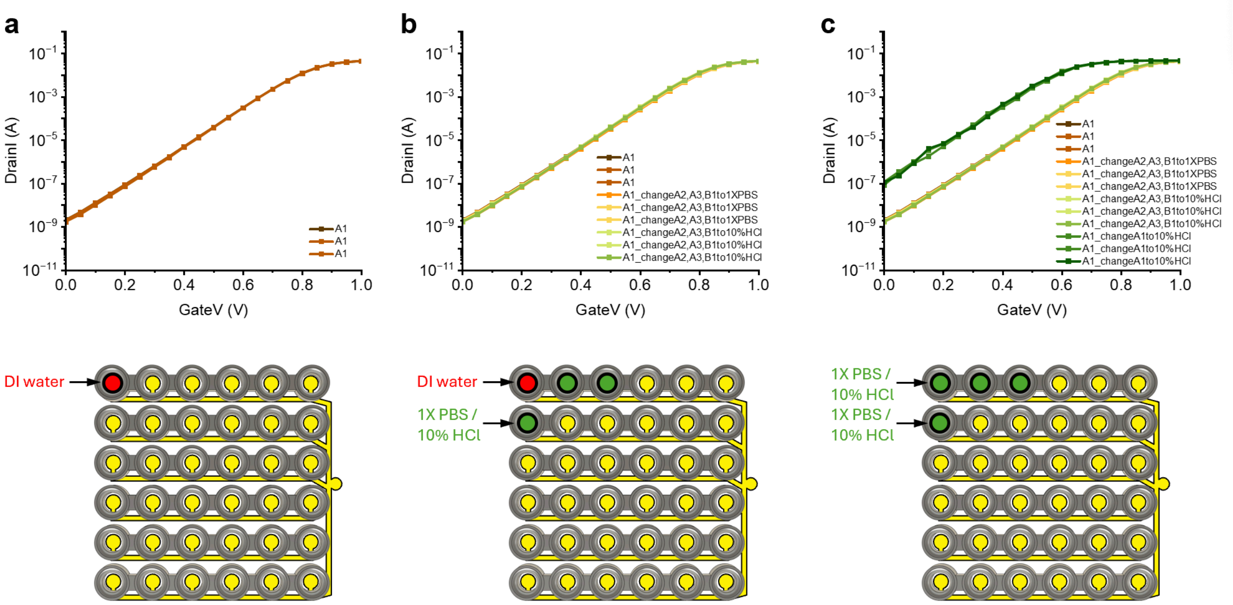


Figure S3. Assessment of cross-talk between wells in the 36-well array. (a) Repeated drain-current versus gate-voltage measurements at well A1 containing deionized (DI) water. (b) Transfer curves at A1 after changing the solutions in neighboring wells A2, A3 and B1 to 1× PBS or 10% HCl, while retaining DI water in A1. (c) Comparison with the response after changing the solution in A1 itself to 10% HCl. The schematics indicate the affected wells. The overlapping curves in (b), compared with the shift upon changing A1 in (c), support limited cross-talk under the tested conditions.

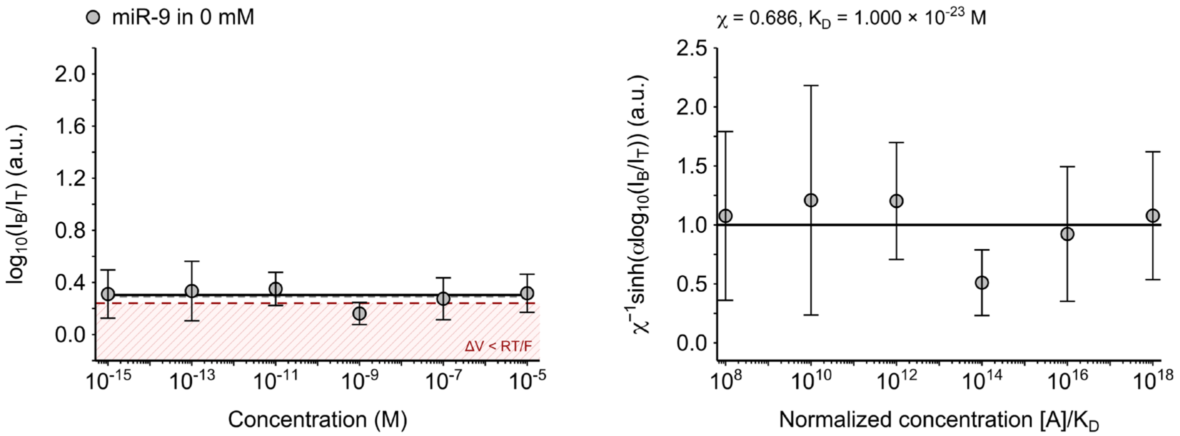


Figure S4. Diagnostic calibration fit for miR-9 without added salt (0 mM). Left: original signal. Right: normalized response.

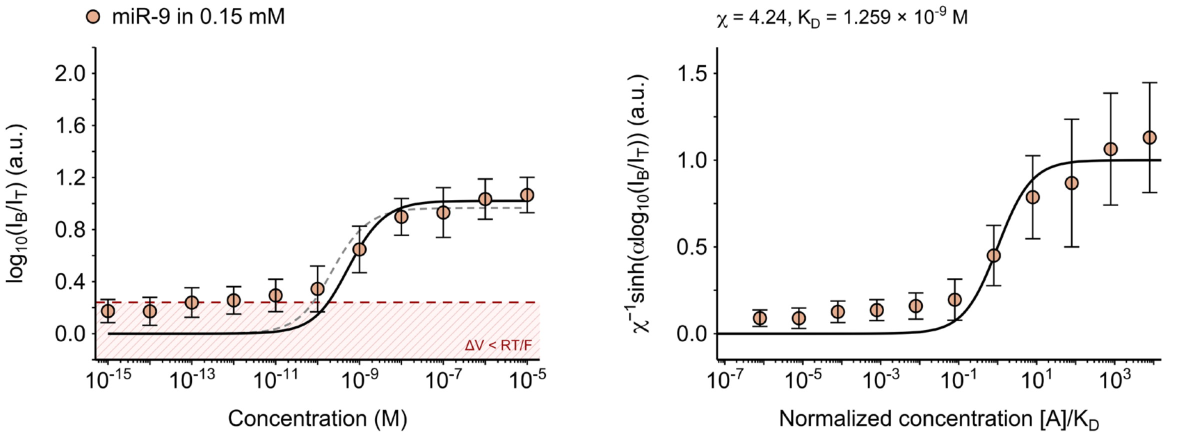


Figure S5. Calibration fits for miR-9 at 0.15 mM ionic strength. Left: original signal. Right: normalized response.

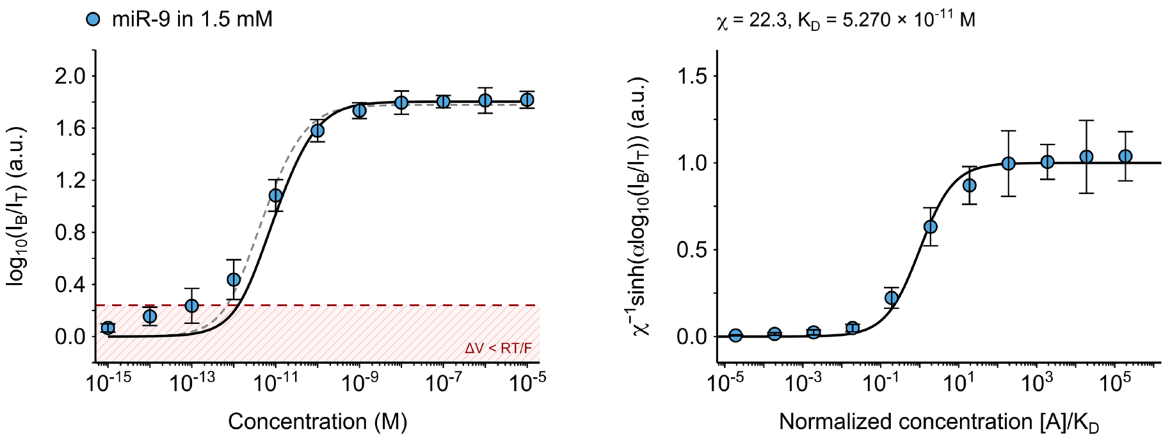


Figure S6. Calibration fits for miR-9 at 1.5 mM ionic strength. Left: original signal. Right: normalized response.

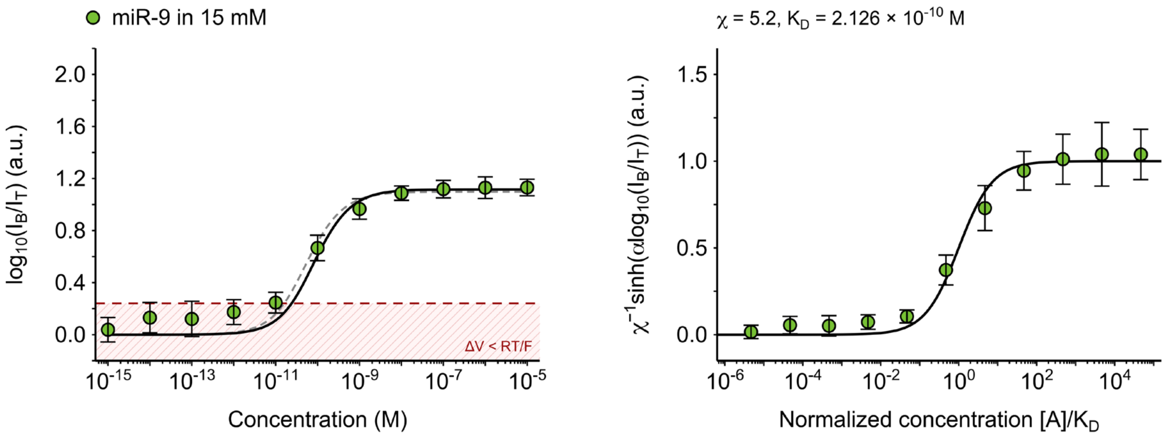


Figure S7. Calibration fits for miR-9 at 15 mM ionic strength. Left: original signal. Right: normalized response.

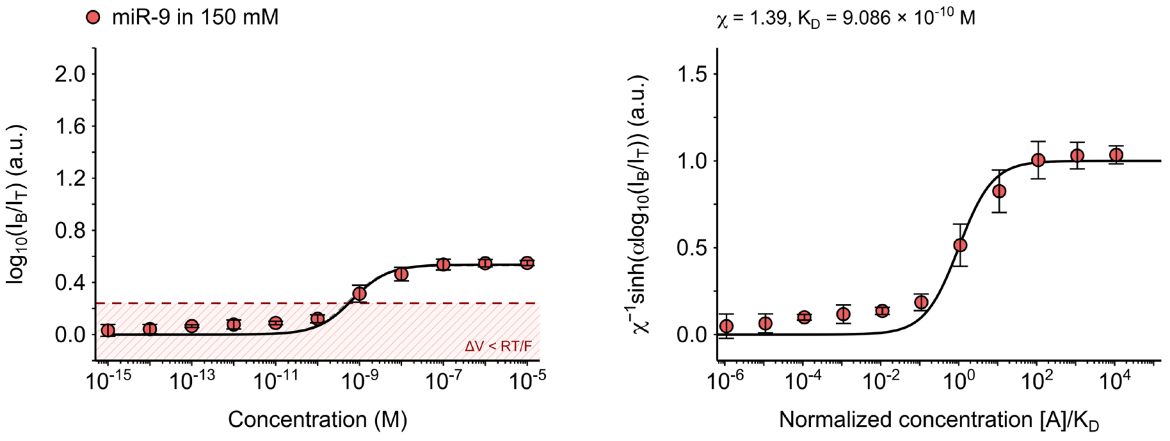


Figure S8. Calibration fits for miR-9 at 150 mM ionic strength. Left: original signal. Right: normalized response.

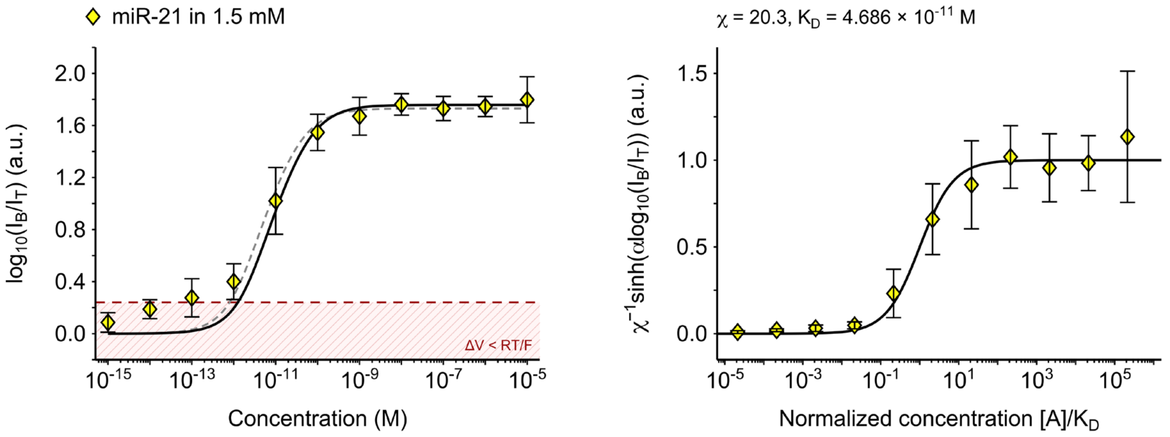


Figure S9. Calibration fits for miR-21 at 1.5 mM ionic strength. Left: original signal. Right: normalized response.

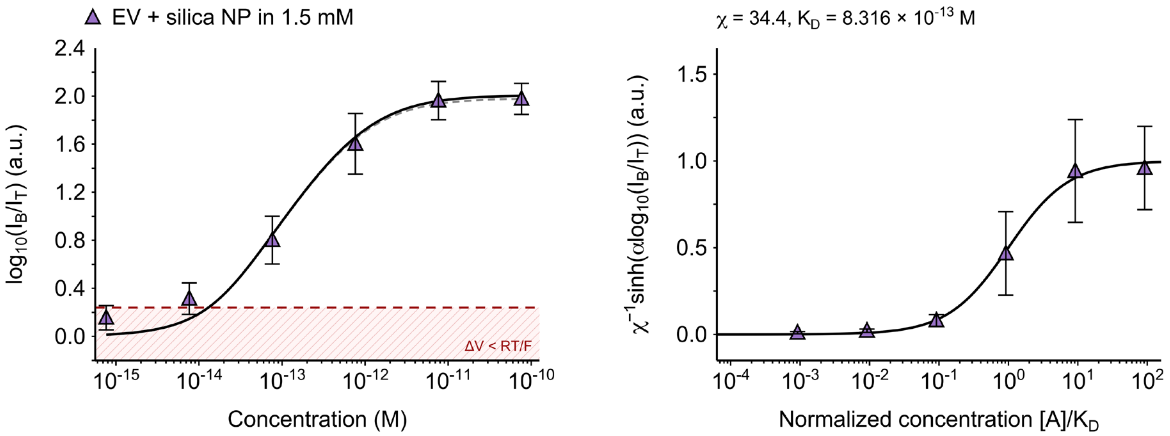


Figure S10. Calibration fits for EVs with silica nanoparticle reporters at 1.5 mM ionic strength. Left: original signal. Right: normalized response.

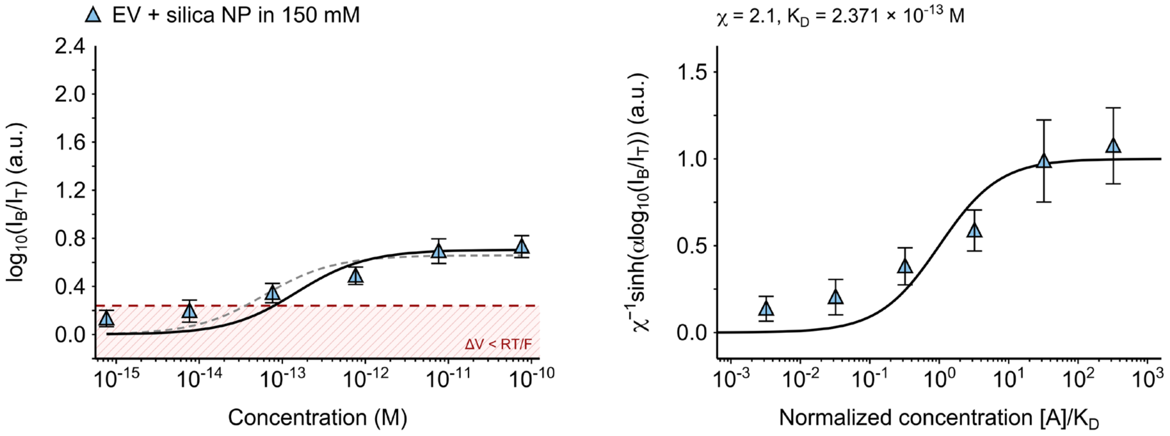


Figure S11. Calibration fits for EVs with silica nanoparticle reporters at 150 mM ionic strength. Left: original signal. Right: normalized response.

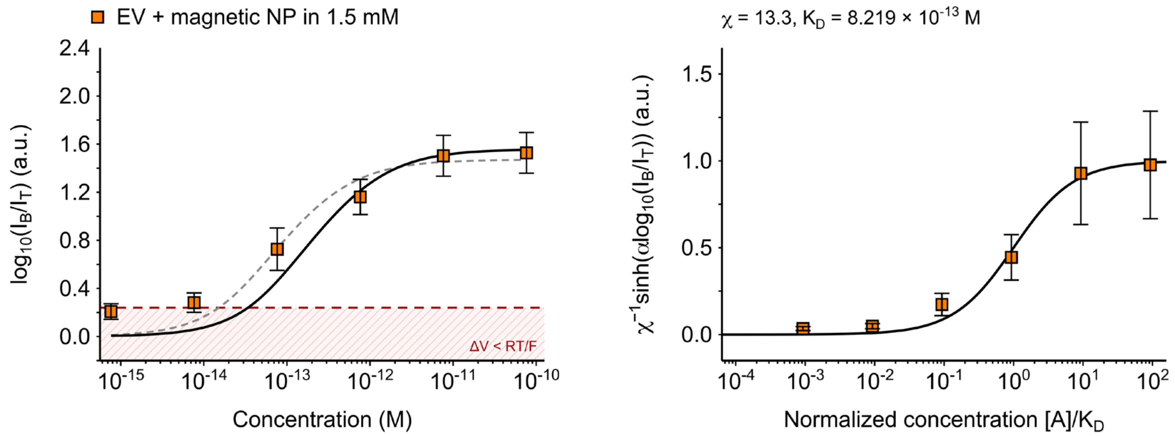


Figure S12. Calibration fits for EVs with magnetic nanoparticle reporters at 1.5 mM ionic strength. Left: original signal. Right: normalized response.

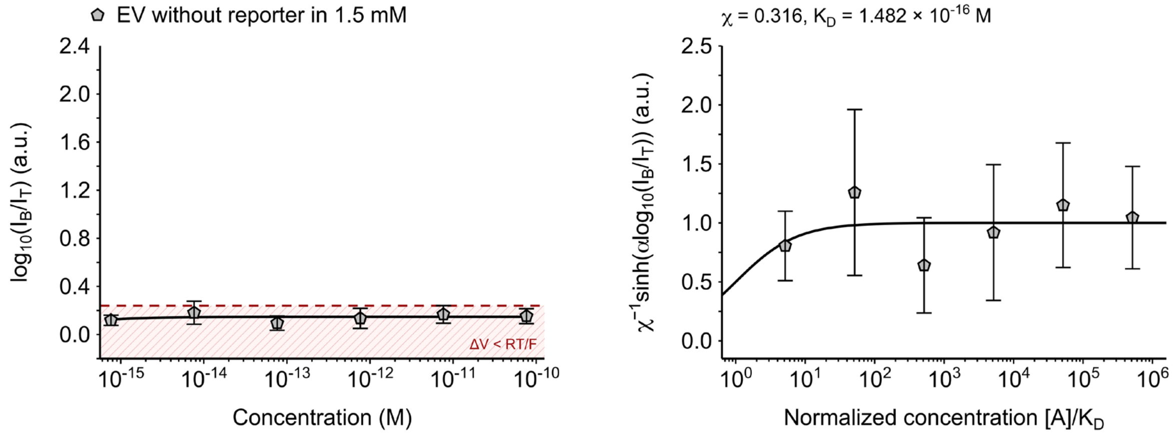


Figure S13. Diagnostic calibration fit for EVs without reporters at 1.5 mM ionic strength. Left: original signal. Right: normalized response.

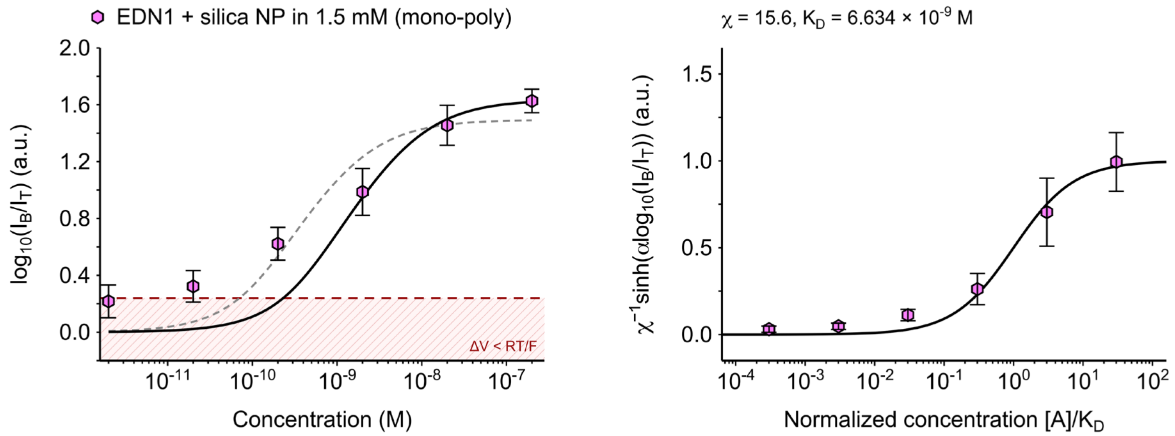


Figure S14. Calibration fits for EDN1 with silica nanoparticle reporters at 1.5 mM ionic strength. Left: original signal. Right: normalized response. Mono-poly denotes monoclonal capture and polyclonal reporter antibodies.

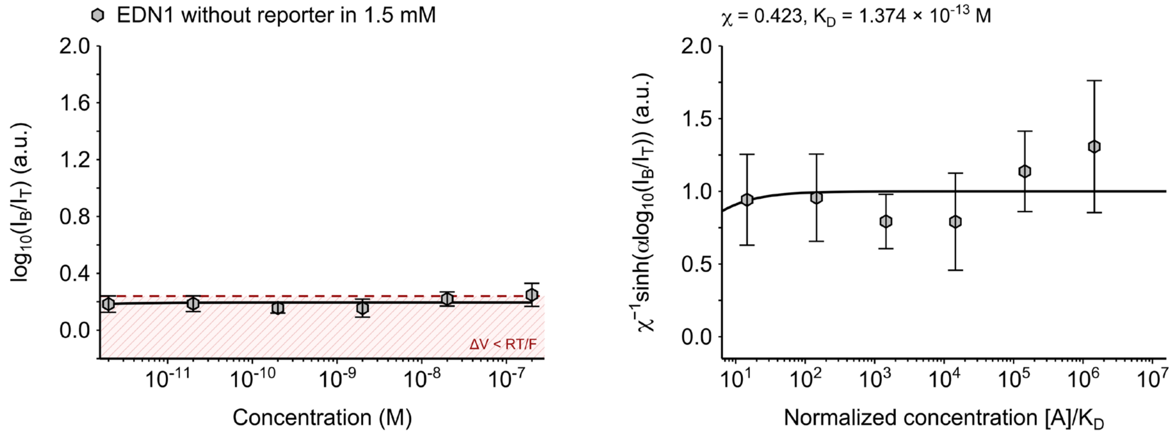


Figure S15. Diagnostic calibration fit for EDN1 without reporters at 1.5 mM ionic strength. Left: original signal. Right: normalized response.

**Table S1. Goodness of fit for the calibration series in Figures S4–S15.**

| Figure | Condition | $R^2$ |
|---|---|---|
| S4† | miR-9, 0 mM | 0.0000 |
| S5 | miR-9, 0.15 mM | 0.7848 |
| S6 | miR-9, 1.5 mM | 0.9534 |
| S7 | miR-9, 15 mM | 0.9482 |
| S8 | miR-9, 150 mM | 0.9438 |
| S9 | miR-21, 1.5 mM | 0.8863 |

| Figure | Condition | $R^2$ |
|---|---|---|
| S10 | EV + silica NP, 1.5 mM | 0.8782 |
| S11 | EV + silica NP, 150 mM | 0.7731 |
| S12 | EV + magnetic NP, 1.5 mM | 0.8586 |
| S13† | EV without reporter, 1.5 mM | 0.0176 |
| S14 | EDN1 + silica NP (mono-poly), 1.5 mM | 0.9239 |
| S15† | EDN1 without reporter, 1.5 mM | 0.0053 |

† Diagnostic controls with little signal response, fitted parameters do not establish reliable binding constants.

Figures S4–S15 compare the output signal $\log_{10}(I_B/I_T)$ with the normalized response $\chi^{-1}\sinh[\alpha \log_{10}(I_B/I_T)]$ of equation (7), where $\alpha \equiv n \ln(10)/2$ and $n = 1.83$. Each well was transformed individually before the normalized means and standard deviations were calculated. The primary fits used unweighted least squares over all transformed well measurements to estimate positive $\chi$ and $K_D$ in $\sinh[\alpha \log_{10}(I_B/I_T)] = \chi\theta([A])$, the fitted form of equation (6). The black curves show the primary fit back-transformed into $\log_{10}(I_B/I_T)$ in the left panels and the ideal Langmuir isotherm $\theta = ([A]/K_D)/(1 + [A]/K_D)$ in the right panels.

The gray dashed curves in the left panels are sensitivity fits obtained by minimizing residuals directly in $\log_{10}(I_B/I_T)$. The red dashed line marks $S = 0.24$, which corresponds to $\Delta V_{analyte} = RT/F$ at $n = 1.83$ This threshold is shown for reference and was not used to exclude measurements from the fits.

The nine responsive conditions have $R^2$ values from 0.7731 to 0.9534 (Table S1). Their normalized plots are consistent with Langmuir-shaped responses, although normalization using fitted parameters is not an independent validation of the model. In contrast, miR-9 at 0 mM and the reporter-free EV and EDN1 controls show little signal response. For miR-9 at 0 mM, $K_D$ reaches the numerical lower bound of $10^{-23}$ M. The parameters and normalized axes for these three controls are therefore diagnostic and do not establish reliable binding constants.
